# Ligand binding remodels protein side chain conformational heterogeneity

**DOI:** 10.1101/2021.09.21.461269

**Authors:** Stephanie A. Wankowicz, Saulo H.P. de Oliveira, Daniel W. Hogan, Henry van den Bedem, James S. Fraser

## Abstract

While protein conformational heterogeneity plays an important role in many aspects of biological function, including ligand binding, its impact has been difficult to quantify. Macromolecular X-ray diffraction is commonly interpreted with a static structure, but it can provide information on both the anharmonic and harmonic contributions to conformational heterogeneity. Here, through multiconformer modeling of time- and space-averaged electron density, we measure conformational heterogeneity of 743 stringently matched pairs of crystallographic datasets that reflect unbound/apo and ligand-bound/holo states. When comparing the conformational heterogeneity of side chains, we observe that when binding site residues become more rigid upon ligand binding, distant residues tend to become more flexible, especially in non-solvent exposed regions. Among ligand properties, we observe increased protein flexibility as the number of hydrogen bonds decrease and relative hydrophobicity increases. Across a series of 13 inhibitor bound structures of CDK2, we find that conformational heterogeneity is correlated with inhibitor features and identify how conformational changes propagate differences in conformational heterogeneity away from the binding site. Collectively, our findings agree with models emerging from NMR studies suggesting that residual side chain entropy can modulate affinity and point to the need to integrate both static conformational changes and conformational heterogeneity in models of ligand binding.

## INTRODUCTION

Ligand binding is essential for many protein functions, including enzyme catalysis, receptor activation, and drug response (Mobley and Dill, 2009). Ligand binding reshapes the protein conformational ensemble between the ligand-bound (holo) and unbound (apo) states, stabilizing some conformations and destabilizing others (Boehr et al., 2009). Despite the dynamic nature of proteins, when comparing structures, often only static conformational changes are considered. However, differences due to ligand binding can range from large, collective movements, such as a loop closure over the binding pocket, to small, local fluctuations of side chains (Gutteridge and Thornton, 2005). Differences in binding affinity and specificity are most often attributed to the enthalpic portion of binding free energy, including visualized interactions between the receptor and ligand. On the other hand, conformational heterogeneity, especially side chain fluctuations, can also contribute energetically to the binding affinity by modulating entropy (Tzeng and Kalodimos, 2012; Wand and Sharp, 2018). A holistic understanding of the origins of binding would ideally explore both enthalpic and entropic energetic contributions to binding affinity (Zhou and Gilson, 2009).

Side chain conformational heterogeneity, including jumps between and variation within rotameric conformations, measured by Nuclear Magnetic Resonance (NMR) relaxation studies has been linked to entropy (Caro et al., 2021; Frederick et al., 2007). In principle, complementary information could be accessed by other structural methods. Structural information from X-ray crystallography or Cryo-electron microscopy (CryoEM), typically produces a single set of structural coordinates. However, the underlying density maps are created from thousands-to-millions of protein molecules, and averaged in both time and space through the crystal lattice or electron microscope particle stack (Cheng et al., 2015; Woldeyes et al., 2014). When averaged in a single density map, conformational heterogeneity across these copies can manifest as “anharmonic disorder”, which can be modeled using multiple alternative conformations, or “harmonic disorder”, which can be modeled by B-factors/atomic displacement parameters (**Figure 1A**). Molecular dynamics experiments have demonstrated that if alternative conformations are not modeled correctly and consistently, then B-factors take on values that are not representative of the underlying conformational heterogeneity (Kuriyan et al., 1986; Kuzmanic et al., 2014). Moreover, B-factors incorporate many effects, including the biases and restraints of the refinement programs, modeling errors, crystal lattice defects, and occupancy changes of atoms. Therefore, consistently modeling X-ray structures as multiconformer models, with alternative side chain and backbone conformations, along with B-factors, may better complement the view emerging from NMR and improve our understanding of the energetics of binding (van den Bedem and Fraser, 2015).

**Figure 1.**
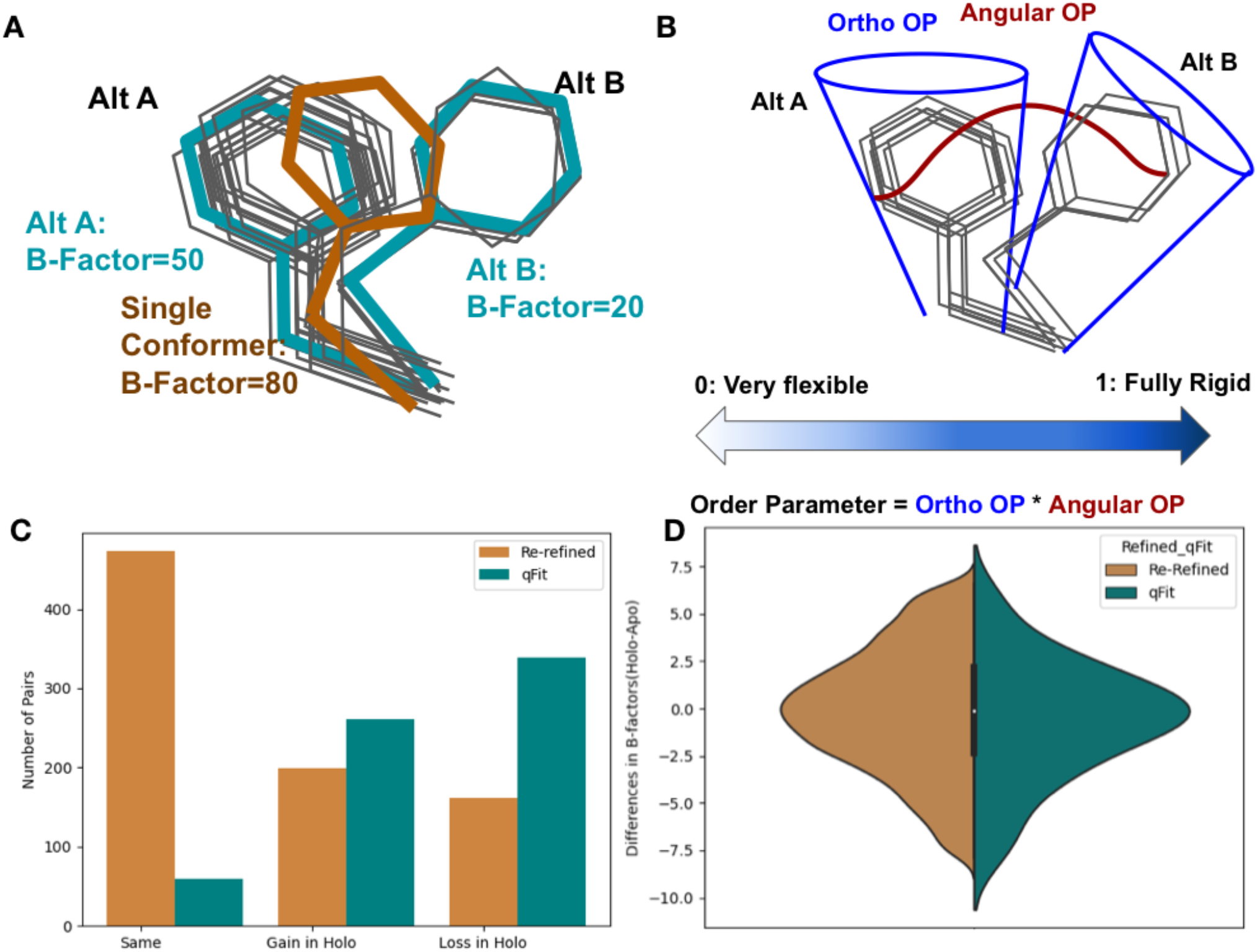
Representing structural data as multiconformer models. (A) The grey outlines represent snapshots of the true underlying ensemble of the phenylalanine residue. The orange stick represents the residue modeled as a single conformer. The teal sticks represent the residue modeled as alternative conformers. The single conformer accounts for all heterogeneity in the B-factor, increasing the B-factor and reducing our ability to determine harmonic versus anharmonic motion. When a residue is modeled using alternative conformers, this heterogeneity is divided between harmonic heterogeneity, captured by the B-factors of each alternative conformation and the anharmonic heterogeneity, captured by spread in coordinates between the alternative conformations. (B) To quantify the conformational heterogeneity of each residue, we used multi-conformer order parameters (Fenwick et al., 2014), which are the products of the ortho order parameter, which captures the harmonic or B-factor portion of each conformation and the angular order parameter, which captures the anharmonic portion or the displacement between alternative conformers. These are multiplied to produce the final order parameter (Methods). (C) The change in the number of alternative conformers (holo-apo) in binding site residues. In the re-refined dataset (orange), the majority structures have the same number of alternative conformers in the binding site, with the second most popular category gaining alternative conformers in the holo structure. In the qfit dataset (teal), the majority of structures lose an alternative conformer in the holo structure, with the second most common category being gaining an alternative conformer. (D) The differences in B-factors (holo-apo) in the re-refined (orange) and qFit (teal) datasets. Overall, there was no significant difference in B-factors between holo and apo structures in both the re-refined and qFit datasets.

Here, we examine how protein side chain conformational heterogeneity changes upon ligand binding by assembling a large, high-quality dataset of matched holo and apo X-ray crystallography structures. To integrate both harmonic and anharmonic disorder, we use a consistent multiconformer modeling procedure, qFit (Riley et al., 2021), and crystallographic order parameters (Fenwick et al., 2014). We test the hypothesis that ligand binding narrows the conformational ensemble, resulting in a decrease in heterogeneity of side chains in the holo structure compared with the apo structure. Our analysis reveals complex patterns of conformational heterogeneity that vary between and within proteins upon ligand binding.

Specifically, in proteins where binding site residues become more rigid upon ligand binding, distant residues tend to become less rigid. This observation suggests that both natural and artificial ligands can modulate the natural composition of the protein conformational heterogeneity across the entire receptor to modulate the free energy of binding.

## RESULTS

### Assembling the dataset

To assess the differences in conformational heterogeneity upon ligand binding, we identified high quality, high resolution (2Å resolution or better) X-ray crystallography datasets from the PDB (Berman et al., 2000). We classified structures as holo if they had a ligand with 10 or more heavy atoms, excluding common crystallographic additives (**Supplementary Table 1, Supplementary Figure 1**). Structures without ligands, excluding common crystallographic additives, were classified as apo (**Supplementary Figure 1A**). We identified holo/apo matched pairs by requiring the same sequence and near-isomorphous crystallographic parameters. Furthermore, we required the resolution difference between holo and apo pairs to be 0.1 Å or less, selecting representative apo structures to minimize the difference in resolution (**Supplementary Figure 1B**). This stringently matched ligand holo-apo dataset contained 1,205 pairs (**Supplementary Table 2**). We also used identical selection criteria to create a control dataset of 293 apo-apo pairs, taken from the set of holo/apo pairs (**Supplementary Table 3**).

### Re-refining and qFit modeling of apo/holo pairs

To minimize biases resulting from different model refinement protocols, we re-refined all structures using the deposited structure factors and *phenix*.*refine* (Liebschner et al., 2019). The majority of structures in our re-refined dataset had less than 2% of residues modeled with alternative conformations, likely reflecting undermodeling of conformational heterogeneity represented in the PDB, based on prior literature (Lang et al., 2010). To more consistently assess conformational heterogeneity, we rebuilt all structures using qFit, an automated multiconformer modeling algorithm (Keedy et al., 2015; Riley et al., 2021) with subsequent refinement using *phenix*.*refine* (Liebschner et al., 2019). While qFit has biases, running all models through a consistent protocol will avoid manual biases that could creep into the holo or apo structures specifically. Additionally, by re-building each model as a multiconformer model, we were able to better distinguish the contributions of harmonic and anharmonic conformational heterogeneity across the structure (**Figure 1A, B**). All models went through additional quality control, removing structures that resulted in large increases in R-free at each refinement step, high clashscores, or large RMSD between the pairs (**Methods**). This procedure resulted in 743 pairs (**Supplementary Figure 2**). Due to apo datasets serving as the reference state for multiple ligand bound structures, our dataset consists of 743 unique holo structures and 432 unique apo structures.

### Properties of the apo/holo pairs

The median resolution across our dataset was 1.6Å with a small trend towards improved (higher) resolution in the apo structure (0.01Å, median improvement (holo-apo); p=3.8×10^−20^, Wilcoxon signed rank test; **Supplementary Figure 3**). Across the dataset, 546 unique ligands were present in the structures, with 134 of these (e.g. NAG, AMP, etc) appearing in multiple structures (**Supplementary Figure 4A**). The median number of ligand heavy atoms was 19, with only 10 very large ligands (>50 heavy atoms, e.g. Atazanavir; **Supplementary Figure 4B**). The proteins in the dataset represent 315 unique Uniprot IDs, with a bias towards enzymes that have been used for model systems for structural biology, including: Endothiopepsin (n=73 pairs), Lysozyme (n=62 pairs), Trypsin (n=48 pairs), and Carbonic Anhydrase 2 (n=46 pairs; **Supplementary Figure 4C, D**).

### Conformational Heterogeneity across the Re-refined and qFit dataset

To determine the differences in conformational heterogeneity upon ligand binding in both the re-refined and qFit models, we assessed four commonly used metrics: the number of alternative conformers, B-factors (atomic displacement parameter), root-mean-square fluctuations (RMSF), and rotamer changes.

### Number of Alternative Conformations

Alternative conformations were modeled at low frequency in the re-refined dataset compared to the qFit modeled structures (1.7% vs. 47.8% of residues). In the re-refined dataset, there is a bias to increased modeling of alternative conformations in the holo dataset (50.5% gain vs. 29.8% loss), whereas more even representation was observed in the qFit dataset (44.3% gain vs. 54.8% loss; **Supplementary Figure 5A**). These results suggest that the trend of increased side chain conformational heterogeneity in PDB deposited structures may have its origin in human bias with more careful human attention to careful model building of binding site residues in holo structures.

We next focused our analysis on binding site residues, defined as any residue with a heavy atom within 5Å of any ligand heavy atom. In the re-refined dataset, 23.9% of the matched pairs had a gain in alternative conformations in the holo model compared to only 19.3% losing an alternative conformer in the holo model, suggesting, counter-intuitively, that ligand binding increases local side chain mobility (**Figure 1C**). However, in the qFit dataset, holo models tend to lose alternative conformations in the binding site residues (39.7% gain vs. 51.5% loss; **Figure 1C**).

### B-factors

Next we explored the harmonic contribution to conformational heterogeneity as modeled by B-factors on a pairwise, residue by residue basis. Across all residues in the re-refined dataset, B-factors were slightly higher in holo models (0.31Å^2^, median difference (holo-apo); p=4.4×10^−208^, Wilcoxon signed rank test; **Supplementary Figure 6A**). In the qFit dataset, similar to the re-refined structures, holo residues had slightly higher B-factors (0.34Å^2^, median difference (holo-apo); p=5.6×10^−264^, Wilcoxon signed rank test; **Supplementary Figure 5B, Supplementary Figure 6**B). Of note, the B-factors in the qFit dataset are slightly smaller than the re-refined dataset (13.41Å^2^ vs. 13.94Å^2^, average B-factors) reflecting the tendency for alternative conformation effects to be modeled as increased B-factors. When examining the binding site residues, there was no significant difference in B-factors between the holo and apo models in both the re-refined (0.01Å^2^, median difference in B-factors; p=0.34, Wilcoxon signed rank test; **Figure 1D**) and qFit datasets (0.06Å^2^, median difference in B-factors; p=0.7, Wilcoxon signed rank test; **Figure 1D, Supplementary Figure 6B**). The lack of change in B-factors close to ligands between the holo and apo models indicate that changes between the holo and apo B-factors are driven by signals distant from the binding site.

### Conformational differences incorporating alternative conformations

Because of the low number of alternative conformers in the re-refined dataset, we only explored the anharmonic differences for side chains between the holo and apo models in the qFit dataset. First, to determine the extent of conformational change of alternative conformations, we compared the rotameric distribution of side chains. Side chain rotamer changes between apo and holo structures have been reported to be very prevalent in single conformer models, with 90% of binding sites having at least one residue changing rotamers upon ligand binding (Gaudreault et al., 2012). To accommodate multiconformer models, we assigned all conformations to distinct rotamers using *phenix*.*rotalyze*. We classified each residue as having “no change” in rotamers if the set of rotamer assignments matched for the holo and apo residue. In binding sites, we also observed that “no change” was the most common outcome for residues (78.6%; **Figure 2A**). In the second largest category, “distinct”, the holo and apo residue shared no rotamer assignments (15.5% of residues; **Figure 2B**).

**Figure 2.**
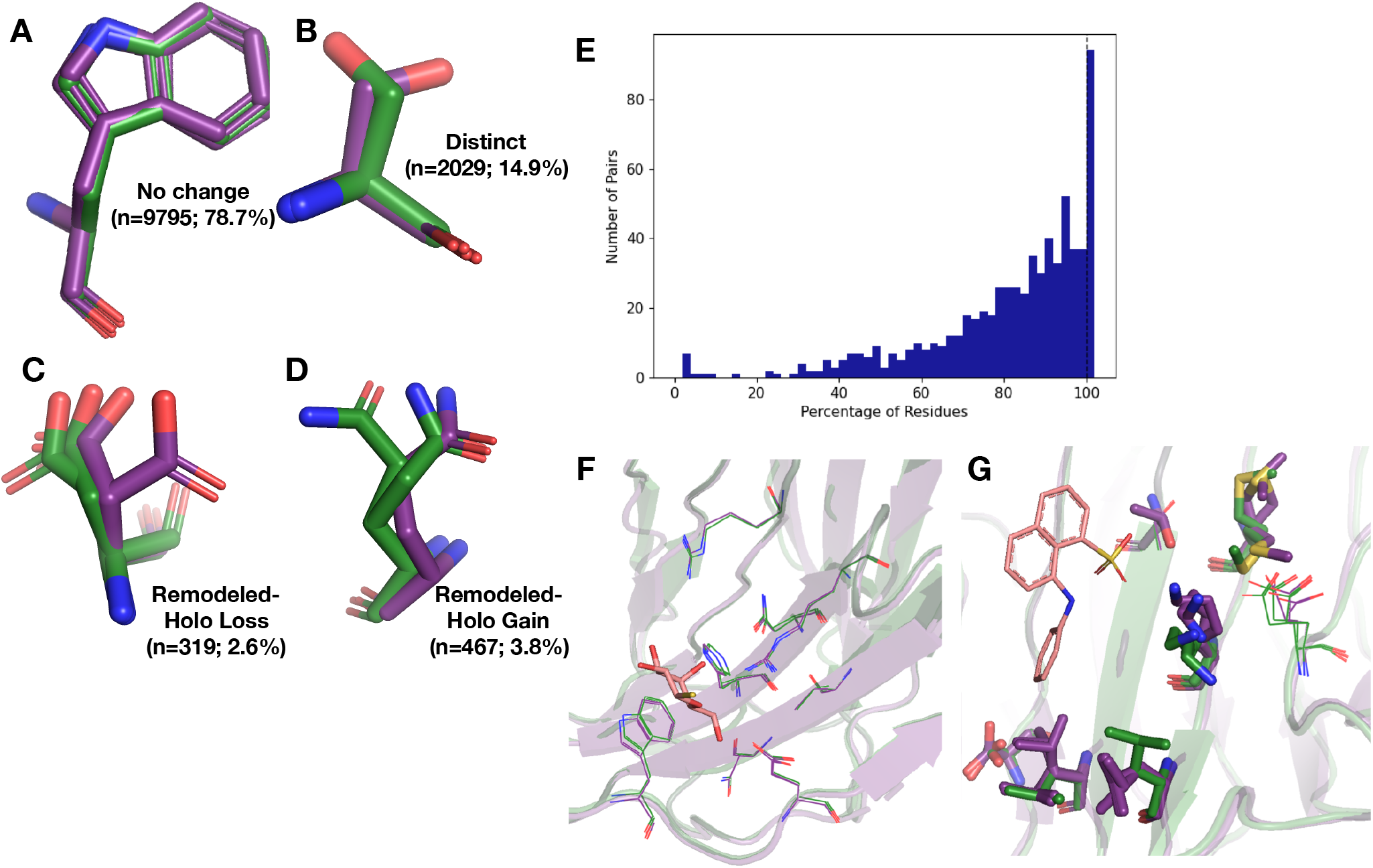
Examples of rotamer changes between apo (purple) and holo (green) binding site residues. (A) Example residues for: ‘no change’ in rotamer status, accounting for 78.7% of binding site residues; (B) “distinct” rotamers, accounting for 14.9% of binding site residues; (C) “remodeled-holo loss”, accounting for 2.6% of binding site residues; and (D) “remodeled-holo gain”, accounting for 3.8% of binding site residues. (E) The percentage of residues in the binding site that have the same rotamer status in the holo and apo structures. The black line highlights the 11% of pairs that had the same rotamer status for all binding site residues. (F) Paired galectin-3 apo (purple; PDB: 5NFC) and holo (green; PDB: 4JC1, ligand: thiodigalactoside) multiconformer models with no changes in rotamer status in any binding site residues. (G) Paired transthyretin apo (purple; PDB: 1ZCR) and holo (green; PDB: 3CFN, ligand: 1-anilino-8-naphthalene) multiconformer models with 6 out of 9 residues with remodeled or different rotamer status in the binding site residues. Residues with rotamer changes are shown as sticks. Residues with the no change in rotamer status are shown as lines.

A more complicated situation occurs when some, but not all, of the rotamer assignments are shared across apo and holo residue. We classified 2.6% of residues as “remodeled - holo loss” (**Figure 2C**) if distinct, additional rotameric conformations were populated in the apo residue only and 3.8% of residues as “remodeled - holo gain” (**Figure 2D**) if distinct, additional rotameric conformations were populated in the holo residue only. These results suggest a counterintuitive interpretation of binding site residues increasing their conformational heterogeneity upon ligand binding. However, a major potential confounder is that holo structures reflect an ensemble average of two compositional states (apo and holo) with alternative conformations representing the apo state at reduced occupancy, which we examined by subsetting the ligands based on relative B-factors (see below). A potential for a third category of remodeling, where both apo and holo residues share at least one conformation and each have at least one additional conformation, did not occur in our dataset.

Across apo-holo pairs there was a large range of the percentages of binding site residues with the same rotamer classification in the pairs (23.2% to 100.0%), indicating that side chain remodeling can be quite variable (**Figure 2E**). We found 11% of binding sites had all residues classified as “same” between pairs, consistent with a previous study that used single conformer models (Gaudreault et al., 2012). As an example of such a “pre-organized” binding site is Galectin-3 bound to thiodigalactoside (PDB: 5NFC, 4JC1; **Figure 2F**). In contrast, 67% of binding site residues have a rotamer status difference in transthyretin (PDB: 1CZR, 3CFN; **Figure 2G**), including a rotamer change in Leu101 to avoid a clash with the ligand.

To compare the magnitude of fluctuations between alternative conformations, we calculated RMSF for all residues. This analysis suggested that, on average, apo residues have slightly greater conformational heterogeneity than holo residues (−0.006Å, mean difference of RMSF(holo-apo); p=3.7×10^−8^, Wilcoxon signed rank test; **Supplementary Figure 6C**). This trend was somewhat stronger in binding site residues (−0.02 Å, mean difference of RMSF(holo-apo); p=4.5×10^−29^, Wilcoxon signed rank test; **Supplementary Figure 6D**). Our RMSF results suggest that, on average, there is a slight decrease in heterogeneity upon ligand binding and that this reduction is most prevalent at residues distant from the binding site.

Collectively, these results do not conform to a simple model. There is a large amount of variability in the response across datasets and the median responses reveal only small biases. Nonetheless, considering those average responses, upon binding a ligand, the RMSF analysis suggests decreases in heterogeneity at the binding site, whereas the rotamer comparison has a slight bias to increased heterogeneity at the binding site, and B-factors only change at distant sites. One interpretation is that heterogeneity is reduced in binding site residues by a small number of anharmonic conformational changes, as observed by the RMSF reduction, paired with an increase in harmonic fluctuations far away, as observed by an increase in the B-factors. However, it is difficult to interpret these changes separately, as conformational heterogeneity is a combination of both harmonic and anharmonic motion and there is potential degeneracy in modeling alternative conformations, even with qFit (van den Bedem et al., 2009). Therefore, we moved to using an integrated measurement of order parameters that can account for these complications (Fenwick et al., 2014).

### Order parameters integrate both harmonic and anharmonic conformational heterogeneity

To integrate the anharmonic fluctuations between alternative conformers with the harmonic fluctuations modeled by B-factors (Kuzmanic et al., 2014), we used a crystallographic order parameter (**Figure 1A**) (Fenwick et al., 2014). While order parameters are traditionally used in NMR or molecular dynamic simulations, they can be calculated for multiconformer X-ray models and, in some cases, show reasonable agreement with solution measures (Fenwick et al., 2014). We focused on the order parameters of the first torsion angle (*χ*1) of every sidechain for all residues except for glycine and proline. Order parameters are measured on a scale of 0 to 1, with 1 representing a fully rigid residue and 0 representing a fully flexible residue. Below, we analyze the differences in normalized order parameters between paired residues (**Methods**).

As an additional control, we compared our apo/holo dataset to a dataset of apo/apo pairs. In examining the differences in order parameters, both in the holo/apo pairs and the apo/apo pairs, there are no large differences in conformational heterogeneity, as indicated by a median difference in order parameters of approximately 0. However, in the holo/apo pairs there is a much wider range of order parameter differences, indicating that ligand binding impacts conformational heterogeneity beyond experimental variability (p=3.4×10^−17^, individual Mann-Whitney U test; **Figure 3A**).

**Figure 3.**
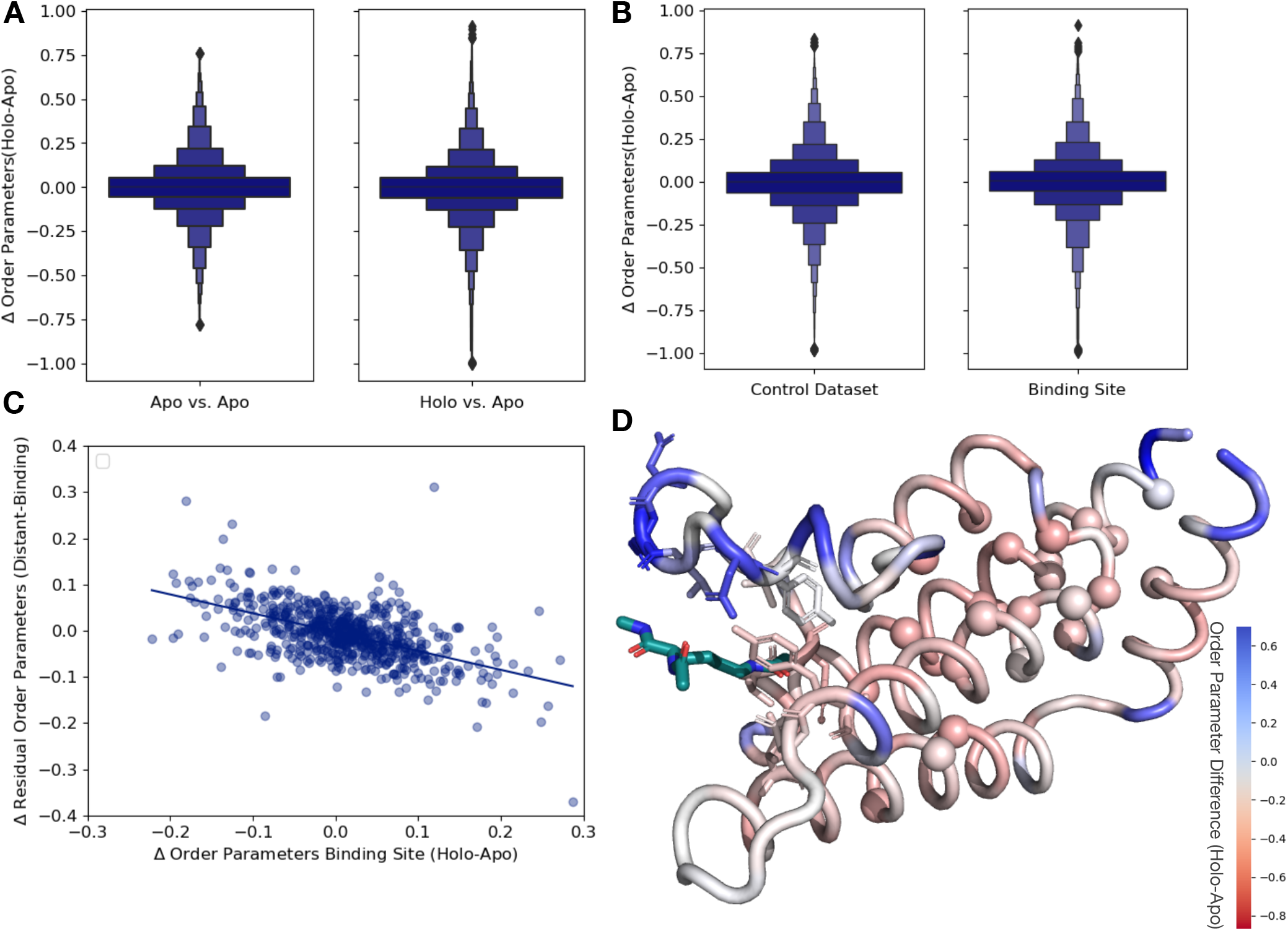
Ligand binding alters conformational heterogeneity patterns. (A) The distribution of order parameter changes is much wider in Holo-Apo pairs compared to Apo-Apo pairs (p=3.4×10^−17^, individual Mann-Whitney U test), however there is no median difference in order parameters upon ligand binding (median difference: 0 for both) indicating that ligands have varying impacts across different proteins. (B) We compare the difference in order parameters in binding site residues of Holo-Apo pairs compared to a control dataset made up of the same number, type, and solvent exposure of amino acids. Comparing the holo/apo structures, on average binding site residues got more rigid upon binding. The median difference in order parameters was 0.03 for the binding site residues, compared to 0 for the control dataset (p=3.4×10^−7^, individual Mann-Whitney U test). (C) The relationship of the difference in order parameters between the holo and apo residues in binding site residues versus the residual order parameter in distant, non-solvent exposed residues. We observed a negative trend (slope=-0.41) indicated that structures that had a loss of heterogeneity in the binding site (right on the x-axis) had a relative gain in heterogeneity in residues far from the binding site that were not solvent exposed (top on the y-axis). (D) We explore this trend on a structure of human ATAD2 bromodomain (PDB: 5A5N). Residues are colored by the differences between the average binding site order parameter minus the order parameter for each residue. Blue residues are less dynamic than the average binding site residue and red residues are more dynamic than the average binding site residue. Binding site residues are represented by sticks and distant, non-solvent exposed alpha carbons are represented by spheres. The ligand ((2S)-2,6-diacetamido-N-methylhexanamide) is colored in teal.

Next, to examine whether different regions of the protein were driving this higher variability, we compared the differences in order parameters among binding site residues, within 5Å of any ligand heavy atom, compared to a control dataset which matched the number of, type and solvent exposure within the protein for each binding site residue. In binding site residues, the holo structures had a slightly, but significantly, increased order parameters, suggesting reduced conformational heterogeneity compared to the control dataset (0.034 vs. 0, median difference (holo-apo) order parameter; p=3.4×10^−7^, individual Mann-Whitney U test; **Figure 3B**). While there is a larger range of responses, this indicated that, in general, binding site residues become more rigid upon ligand binding.

### Spatial distribution of conformational heterogeneity changes

Based on the large range of order parameter differences we observed across the protein, along with the decrease in heterogeneity localized to binding site residues, we next explored the relationship between changes in heterogeneity in binding site residues and the rest of the protein. The difference in order parameters between the holo and apo models were correlated in both the binding site and distant residues (**Supplementary Figure 7A**), indicating that ligand binding generally caused global changes to flexibility. Given the average rigidification of the binding site residues (**Figure 3B**), these results predict a general trend of decreased conformational heterogeneity in the ligand binding site would be associated with a relative increase in conformational heterogeneity at distant sites in the protein. This pattern suggests that the residual change in heterogeneity (the difference between the average order parameter of the distant residues and the average order parameter of the binding site residues) should be inversely related to the change in the binding site residues: more rigidified binding sites will have more flexible than expected distant sites, and vice versa. Indeed, on a protein-by-protein basis, the relationship between binding site residues and residual changes at distant sites follows this trend (**Supplementary Figure 7B**). Consistent with studies suggesting significant residual conformational heterogeneity in folded buried residues (Wong and Daggett, 1998) and the potential for those buried residues to change heterogeneity upon ligand binding (Moorman et al., 2012), this trend is even stronger in residues that were more than 10Å away from any heavy atom in the ligand and less than 20% solvent exposed (slope=-0.41, r^2^=0.46; p=5.1×10^−50^, two-sided t-test; **Figure 3C**). This indicates that proteins that lose conformational heterogeneity in the binding site are associated with a relative increase in conformational heterogeneity in distant, non-solvent exposed residues.

There are three likely origins of this effect. First, this may reflect a feature of the distribution of order parameters around the mean value within each protein. Second, this may reflect a topological feature of protein packing, whereby packing optimization of certain areas of a protein decreases the optimization of other parts of the protein (Bromberg and Dill, 1994). Third, this may reflect the stabilization of certain conformations in a ligand bound protein. As a control for these effects, we compared the residual order parameter differences between the buried, non-solvent exposed residues and the binding site residues in apo-apo pairs. Globally the trends were similar, but weaker in both correlation and magnitude (slope=-0.28, r^2^=0.20; **Supplementary Figure 7C**). Therefore, we interpret the trend we observe as mainly based on protein topology, specifically that proteins have areas where there are less efficiently packed alternative conformers, likely to enable entropic compensation across the protein during various functions, including ligand binding. We interpret that the stronger signal we observed in the holo-apo dataset is due to the ligand perturbation, which is also reflected in the median rigidification of binding site residues (**Figure 3B**). We hypothesize that we are observing this innate protein property being used, specifically optimizing the binding site residues to bind a ligand, while decreasing the optimization elsewhere in the protein.

As an example to visualize this trend, we mapped the change in order parameters onto the structure of the human ATAD2 bromodomain (PDB ID:5A5N). In ATAD2, the binding site residues rigidify upon ligand binding whereas the majority of distant residues are more heterogeneous compared to the binding site residues (**Figure 3D**). Specifically, this difference is greatest between binding residues and non-solvent exposed residues. However, as in the global analysis, this example demonstrates there is a large range of changes in binding site order parameters, consistent with NMR examples that show a heterogeneous response both close to and far from ligands (Caro et al., 2021; Moorman et al., 2012).

### Ligand properties influence conformational heterogeneity

Next we investigated how ligand properties impact the conformational heterogeneity of binding site residues. For ligand properties dictated by the size of the ligand (number of rotatable bonds and number of hydrogen bonds) we normalized these metrics by the molecular weight of the ligand. For each property, we compared the highest and lowest quartiles by both the absolute order parameters of the holo structure and the order parameters differences between holo and apo pairs. No significant associations existed when comparing the differences between holo and apo order parameters, but the characteristics of the holo binding site and the rotamer changes were correlated with ligand properties in several cases.

We hypothesized that ligand properties associated with increase ligand dynamics, including more rotatable bonds, higher lipophilicity (logP), fewer hydrogen bonds, and more heavy atoms would be associated with increased conformational heterogeneity (an increase in absolute order parameters or a smaller difference between the apo and holo order parameters; Wicker and Cooper, 2015). While molecules with fewer rotatable bonds (lower quartile: <2 (n=134) vs. upper quartile: >6 (n=134)) were indeed associated with more rigid binding sites (lower quartile: 0.83 vs. upper quartile: 0.81, individual Mann Whitney U test), this was not significant. However, higher numbers of rotatable bonds were associated with a lower number of same rotamers between the apo and holo binding site residues (88% vs. 80%, percentage same rotamer; p=6.0×10^−6^, individual Mann Whitney U test; **Supplementary Figure 8A**). Increased lipophilicity (logP, upper quartile: <0.04 (n=134) vs. lower quartile: >2.69 (n=134)), was significantly associated with a more flexible binding site (0.84 vs. 0.79, median order parameters; p=7.5×10^−6^, individual Mann Whitney U test; **Figure 4A**). Previous studies have indicated that increased lipophilicity generates more nonspecific binding interactions (Olsson et al., 2008). Larger compounds (upper quartile: >26 heavy atoms (n=134) vs. lower quartile: <13 heavy atoms (n=134)) are also associated with more flexible binding sites (0.79 vs. 0.83, median order parameter; p=0.0001, individual Mann Whitney U test; **Figure 4B**). Large compounds, thus a larger ligand surface area, are associated with more nonspecific binding interactions, which is compatible with increased protein conformational heterogeneity. Finally, more total hydrogen bonds per heavy atom(upper quartile: >0.47 (n=134) vs. lower quartile:<0.25 (n=134)) are associated with more rigid binding sites (0.84 vs. 0.79, median order parameter; p=5.9×10^−5^, individual Mann Whitney U test; **Figure 4C**). This trend holds even when examining hydrogen bond donors or acceptors separately.

**Figure 4.**
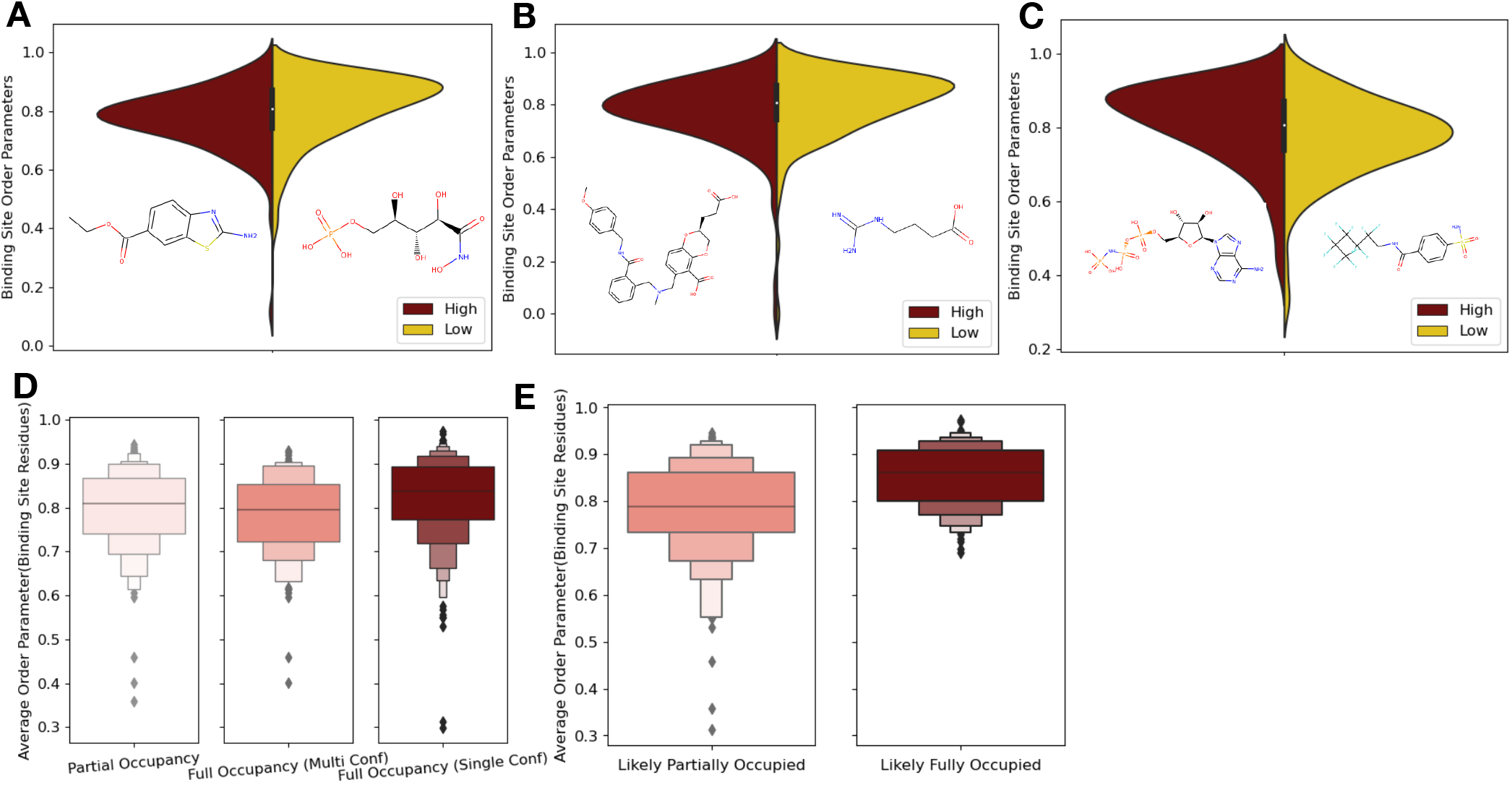
Ligand properties impact binding site order parameters. (A) Ligands with higher logP value (maroon), indicative of more greasy or hydrophobic ligands, versus ligands with a lower logP value (gold), had lower in order parameters in the binding site residues (0.78 vs. 0.84, median order parameter; p=7.5×10^−6^, independent Mann Whitney U test) [Example ligands: low logP: 5-phospho-d-arabinohyroamic acid, high logP: ethyl 2-amino-1,3-benzothiazole-6-carboxylate]. (B) Ligands with relatively higher molecular weight (maroon) had higher order parameters compared to those with lower molecular weight(gold; 0.79 vs. 0.83, median order parameter; p=0.0001, independent Mann Whitney U test). [Example ligands: High number of heavy atoms: (2S)-2-(3-hydroxy-3-oxopropyl)-6-[[[2-[(4-methoxyphenyl)methylcarbamoyl]phenyl]methyl-methyl-amino]methyl]-2,3-dihydro-1,4-benzodioxine-5-carboxylic acid, low number of heavy atoms: 4-carbamimidamidobutanoic acid]. (C)Ligands with relatively higher hydrogen bonds per heavy atom (maroon) had higher order parameters compared to those with lower molecular weight (gold; 0.84 vs. 0.79, median order parameter; p=5.9×10^−5^, independent Mann Whitney U test) [example ligands: low hydrogen bond: 4-sulfamoyl-N-(2,2,3,3,4,4,5,5,6,6,6-undecafluorohexyl) benzamide, high hydrogen bond: phosphoaminophosphonic acid-adenylate ester]. (D) Binding site order parameters were lower in ligands with partial occupancy (light pink; 0.79, median order parameter) and mutliconformer ligands adding to full occupancy (salmon; 0.80, median order parameter), compared to single conformer ligands with full occupancy (dark red; 0.83, median order parameter; p=4.9×10^−8^, independent Mann Whitney U test). (E) In fully occupied ligands, ligands in the top quartile of ligand B-factors, controlled for by the mean alpha carbon B-factor, had lower binding site order parameters (salmon; 0.79, median order parameter) compared to ligands in the bottom quartile (dark red; 0.85, median order parameter; p=1.6×10^−11^, independent Mann Whitney U test).

From these results, an intuitive general picture emerges where more specific interactions, such as hydrogen bonds, are correlated with more rigid binding site residues, whereas the more non-specific interactions are correlated with more flexible binding site residues. There is also a wide range of deviation from this general picture, likely reflecting that natural and artificial optimization of ligands is based on free energy, not any specific thermodynamic component or interaction type. These trends emphasize the need to monitor both the impacts of ligands on specific interactions with the protein along with conformational heterogeneity of the protein. Additionally, these results suggest that specific interactions can be tuned to rigidify a binding site. Paired with our findings of the relationship between order parameters in binding site and distant residues, ligand impacts are likely propagated throughout the protein. Ligands with more specific interactions, thus a less flexible binding site, will likely have a corresponding increase in conformational heterogeneity far away from the binding site.

### Reduced ligand occupancy and conformational heterogeneity

One potential confounder for quantifying the change in conformational heterogeneity of binding site residues is that the ligands may not be fully occupied in the crystal. There were 193 structures with ligands with alternative conformations or partially occupied ligands in our datasets (**Figure 4D**). Of these 193, 125 ligands had less than full occupancy, whereas 68 had alternative conformations that amounted to full occupancy. The vast majority of ligands (n=425) were modeled originally with full occupancy. Fully occupied ligands were associated with more rigid binding sites than partially occupied ligands or ligands with alternative conformers (0.84 vs. 0.79, mean order parameters of binding site residues; p=2.9×10^−7^, individual Mann Whitney U test; **Figure 4D**). There was no difference observed between the partially occupied ligands and ligands with alternative conformers (p=0.15, individual Mann Whitney U test; **Supplementary Figure 8B**). We also explored if partially occupied ligands were associated with more rotamer changes between holo and apo pairs, but no significant difference existed (80% vs. 85%, median percentage of the same rotamer;**Supplementary Figure 8B**).

While the scattering contributions of B-factor and occupancy changes are subtle (but distinct), most models likely include true occupancy changes as elevated B-factors. We observed a wide range of average ligand B-factors and, as expected, a lack of correlation between the ligand B-factors and ligand occupancy (Bhat, 1989; Carugo, 1999; van Zundert et al., 2018). As a proxy for likely partially occupied ligands, we normalized the ligand B-factor by the mean C-alpha B-factor to identify ligands with higher B-factors than expected (**Supplementary Figure 8C**). We examined the outer two quartiles of the normalized ligand B-Factors (>0.016 vs. <0.005, median normalized B-factor). In these “likely partially occupied” ligands, we observed greater conformational heterogeneity (0.86 vs 0.80, mean order parameter; p=1.6×10^−11^; individual Mann Whitney U test, **Figure 4E**). In structures with modeled partially occupied ligands and likely partially occupied ligands, we learned that binding site residues tend to have more apparent conformational heterogeneity, likely due a combination of compositional and conformational heterogeneity.

### Conformational heterogeneity for multiple ligands to CDK2

To better understand our findings in the context of multiple, diverse ligands binding to a single receptor, we examined Cyclin-dependent Kinase 2 (CDK2), a cyclin kinase family that regulates the G1 to S transition in the cell cycle. Our dataset contains 13 protein-inhibitor complexes, including both type I and type II inhibitors, all of which share the same apo model (PDB ID: 1PW2). We hierarchically clustered the residues and ligands by difference in order parameters between the holo and apo models, identifying three distinct clusters of residues. The first cluster (blue, **Figure 5A,B**), consisting of 13 residues, are rigidified upon ligand binding. This cluster included residues scattered throughout both the N- and C-lobes of CDK2 that rigidified upon ligand binding. This dispersed pattern is similar to the trend of rigidification that is observed by NMR in PKA upon substrate binding (Kim et al., 2017). Two notable residues in this cluster, Glu127 and Val18, contact the inhibitors. Upon ligand binding, Val18 transitions from multiple conformers to a single conformation. Glu127 has a similar conformation in the apo and type II structures of two distinct alternative side chain rotamers, whereas in the type I inhibitor structure, the alternative conformers cluster around the same rotamer (**Supplementary Figure 9A**,**B**).

**Figure 5.**
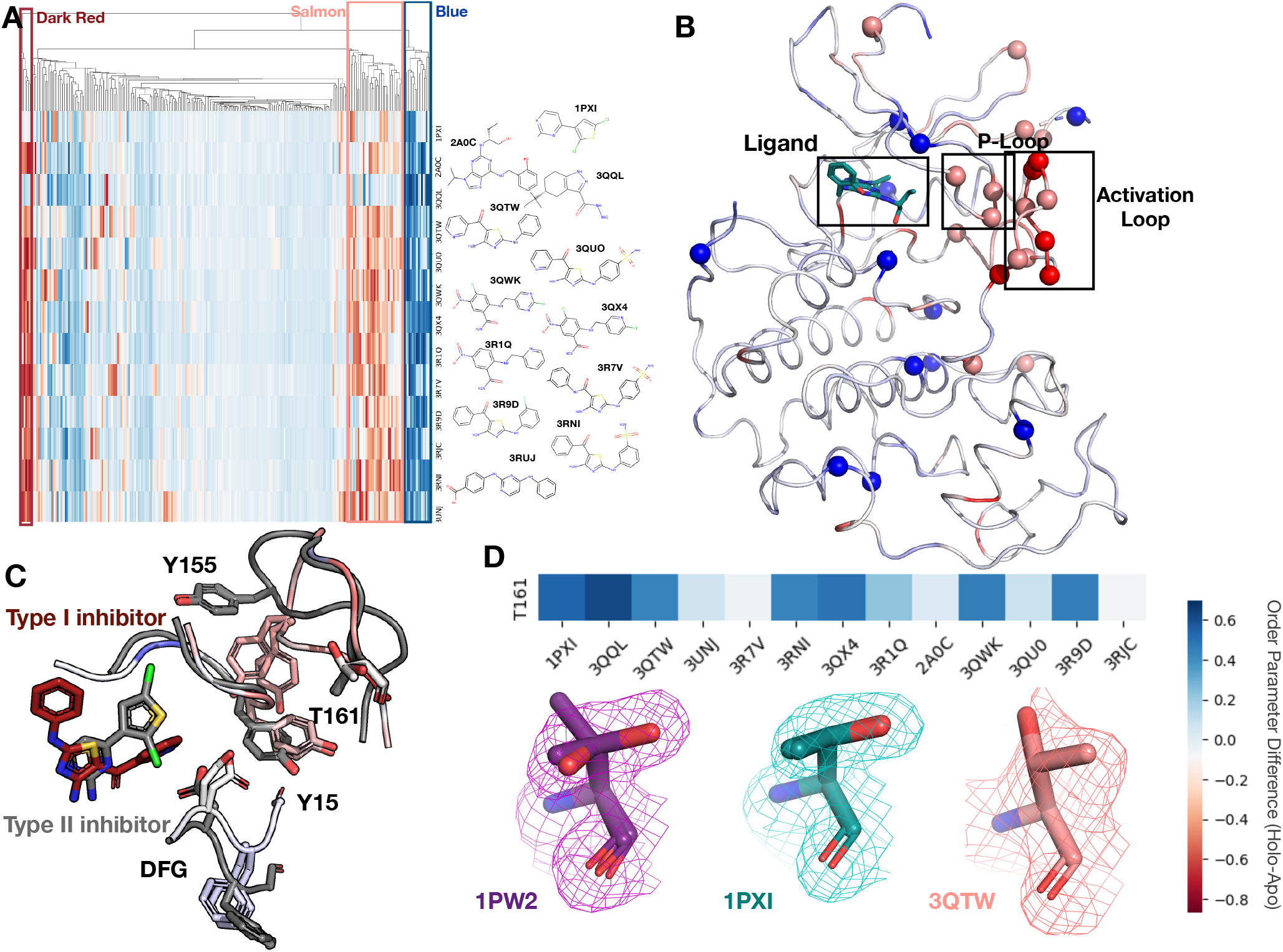
Conformational change and heterogeneity in CDK2. (A) The clustermap of all residues in the 13 CDK2 protein/ligand pairs. Red values indicate a negative difference (holo-apo) in order parameters, indicating that the holo structures have more heterogeneity compared to the apo. Blue values indicate positive differences, indicating that the apo structures have more heterogeneity compared to the holo. We highlighted three important clusters, the left red cluster, middle salmon cluster, and right blue cluster. (B) A representative structure (PDB: 3QTW) is shown with each residue colored by the difference in order parameter, corresponding to the same coloring scheme as the clustermap. The three distinct clusters (dark red, salmon, blue) are shown in spheres. (C) Many of the key differences between type I inhibitor(PDB: 3QTW) and type II inhibitor(PDB: 1PXI) are located in the DFG motif, p-loop, and activation loop. The type II inhibitor structure is colored in grey and the type I inhibitor is colored as the difference in order parameters between the type I inhibitor and type II inhibitor structures. Red signifies a more dynamic region in the type I inhibitor structure, blue signifies a less dynamic region in the type I inhibitor structure. Changes in the DFG motif, propagates changes, both structural and in dynamics, in the p-loop (highlighted by Tyr15), which progates even larger changes in the activation loop between the two inhibitors, including changes in conformation of Thr161, the phosphorylation site of CDK2. (D)Threonine 161, the phosphorylation site for CDK2. We looked at the supporting density for specific residues between the apo (PDB: 1PW2, purple), type II (PDB: 1PXI, teal), and type I (PDB: 3QTW, salmon) inhibitors. 2Fo-Fc electron density is shown at 1 sigma. The apo structure has multiple conformations, whereas the both holo models only have one, but in different rotamer states

The second cluster (salmon, **Figure 5A**,**B**), consists of 14 residues that increase flexibility upon ligand binding. The majority of these residues connect the p-loop and the activation loop (**Figure 5B**). The electron density is very weak for many of these residues in most of the holo structures, driving their modeling in multiple conformations and elevated B-factors (**Supplementary Figure 9D**). The third cluster (dark red, **Figure 5A**,**B**) is comprised of five residues that became more flexible in all, but two holo datasets, which are the only type II inhibitors in the dataset. These were all located on the activation loop of the kinase (**Figure 6D**). As type II inhibitors, the two molecules [PDB: 1PXI (ligand: CK1) and PDB: 3QQL (ligand: X03)] bind the DFG out conformation present in the apo dataset (PDB: 1PW2) and do not have as drastic of a rigidifying effect as the type I inhibitors. Notably, these two inhibitors were also smaller than the type I inhibitors and the reduced contacts may also drive some of this effect.

The multiconformer models also provide a structural rationale for these changes. The differences in DFG conformation change the contacts with the P-loop, which allow for greater side chain flexibility in the “up” form compatible with type I inhibitors. The interface between the P-loop and the activation loop is weaker and residues such as Tyr155 adopt multiple conformations. At the base of the activation loop, Thr161, a critical phosphorylation site, changes conformation, with a rigidifying effect common to both type I and II inhibitors (**Figure 5C**). The conformation of Thr161 found in the type II inhibitors overlap, with one of the conformations populated in the multiconformer apo model. In contrast, the type I inhibitors adopt a distinct conformation. This case study highlights how modeling information present in the density can reveal changes beyond those in single conformer structures.

## DISCUSSION

By creating a large dataset of stringent matched pairs of apo and holo multiconformer models, we identified a pattern of conformational heterogeneity consistent with smaller scale studies of individual proteins (Caro et al., 2021). In general, we found that binding site residues tend to become more rigid upon ligand binding. However there was a large range of effects, including many sites becoming more flexible when bound to a ligand. The trends suggest that disorder-order transitions between binding site residues and distant residues are common and potentially a selected property of many proteins (Kim et al., 2017). Specifically, our data suggests that some of the entropy lost by the rigidification incurred by binding site residues upon ligand binding can be compensated with an increase in disorder in distant residues. This finding generalizes the phenomenon has been observed in single protein analyses with X-ray, NMR, and MD simulation (Gohlke et al., 2004; Moorman et al., 2012; Wang et al., 2019). Both theoretical and experimental analyses suggest that the relationship between local packing optimization and small voids that permit alternative conformations will be key to predictably mapping this relationship (Bromberg and Dill, 1994; Caro et al., 2021). Using temperature or pressure as perturbations during X-ray data collection can help to further map the connection between packing “quality” and side chain conformational heterogeneity in greater detail (Caro et al., 2021). While NMR order parameter studies only take into account movement that is shorter than the tumbling time for the protein (Hoffmann et al., 2021; Gangé et al., 1998), our results are insensitive to timescale. In addition, it is quite likely that our use of cyro-cooled structures causes an underestimate of the heterogeneity occurring in these datasets (Fraser et al., 2011).

We observed a complex interplay between conformational change and dynamics in our analysis of 13 inhibitor-bound datasets of the kinase CDK2, in the same crystal form and space group. The ability to explore one protein with multiple ligands highlights the utility of crystal systems amenable to isomorphous soaking or co-crystallization (Steuber et al., 2006). We identified differences in conformational heterogeneity between type I and type II inhibitors that can be classified along with well-known changes, such as differences in the DFG motif. Tuning distal site dynamics may be a viable strategy for modulating the affinity of kinase inhibitors and affect the pattern of protein-protein interactions on distal surfaces, which is of critical importance in CDK inhibitor development (Jhaveri et al., 2021; Wood and Endicott, 2018).

We note that our work is not sensitive to many facets of the complex changes associated with ligand binding (Mobley and Dill, 2009). Our stringent resolution matching criterion may also render us blind to the most severe effects on conformational heterogeneity, whereby ligand binding causes a more widespread change leading to a loss or gain of diffraction power. In addition, water molecules play an important role in ligand binding, both in the release of ordered water molecules contributing to binding through entropy and in forming specific interactions (Breiten et al., 2013; Verteramo et al., 2019). Additionally, ligand conformational heterogeneity has been highlighted by several recent studies (Caldararu et al., 2021; Jain et al., 2020; van Zundert et al., 2018). Another caveat in our analysis is the limitations of qFit modeling for modeling extensive backbone heterogeneity into weak electron density. Ensemble modeling methods, which leverage molecular dynamics for sampling and use a different model representation may be more appropriate for examining these systems (Burnley et al., 2012; Eshun-Wilson et al., 2019). Future work, integrating the conformational heterogeneity of the protein, ligand, and water molecules will create better predictions and explanations of the energetics of binding. In addition, this would allow us to interpret the impact of specific interactions and alterations on both the entropy and enthalpy of all components of the system.

Our study, as well as previous NMR studies (Caro et al., 2017; Frederick et al., 2007), only leverage a limited set of side chain dihedral angles. However, comparisons with molecular dynamics simulations suggest that small sets of side chain dihedrals alone may be representative of the overall changes in dynamics of the system (Wand and Sharp, 2018, Fleck et al., 2018). What is the thermodynamic impact of restricting side chain conformational heterogeneity? Protein folding studies and theory indicate that restricting the rotamer of even a single side chain can incur an entropic penalty of binding of ∼0.5kcal/mol (Doig and Sternberg, 1995). While we observe many such restrictions in binding sites due to specific interactions with ligands, our data point to corresponding changes away from the binding site that help balance this cost. Overall, the median increase in rigidity we observe in binding site residues (0.03 order parameter increase) would create an energetic penalty of approximately ∼0.1-0.5kcal/mol (Wand and Sharp, 2018), with outliers having even larger thermodynamic consequences. Given the constraints of maintaining a folded conformational ensemble upon ligand binding, it is likely that ligand binding generally acts to restrict degrees of freedom locally and that protein topological constraints favor increased motion in distal regions (Bromberg and Dill, 1994). This overall effect likely manifests because optimizing affinity is desirable for medicinal chemistry and for the selective pressures experienced by many proteins. Such optimization is insensitive as to whether the free energy is driven enthalpically or entropically. However, given the attention paid to designing and optimizing enthalpic interactions, there is likely unleveraged potential in optimizing the entropic component as well. As more sophisticated models of conformational heterogeneity are created and validated (Rosenbaum et al., 2021) the strategy of rationally tuning conformational heterogeneity to improve binding affinity may be an attainable design strategy.

## METHODS

### Dataset

Our dataset was compiled using a snapshot of the PDB(Berman, 2002) in September 2019, containing 156,187 structures. We then selected structures that had a resolution better or equal to 2Å (n= 64,557). We also excluded any structure that contained nucleic acids (n=2,280) or covalently bound ligands (n=1,030). We identified holo structures(n=30,530), defined as those that contained at least one ligand, defined as any HETATM residue with 10 or more heavy atoms, excluding common crystallographic additives.

To create apo/holo pairs, we took each holo structure and compared it to each potential apo structure (n=30,717), defined as structures without a ligand bound. A pair was defined according to the following criteria:

- same space group
- exact sequence or exact sequence after removing the first or last five base pairs of either structure
- a resolution difference between the two structures less than 0.1Å
- dimensions of unit cells do not differ by more than 1Å
- angles of the unit cells do not differ by more than 1 degree

This gave us 15,214 pairs. We then subsetted this list down to provide only one apo structure per holo structure, based on the smallest resolution difference. This produced a final pair set of 1,205 with 1,143 unique structures (**Supplementary Table 2**).

We also created a pairset with 458 unique apo/apo pairs using the same criteria as the apo/holo pairset (**Supplementary Table 3**).

### Refinement

We re-refined all structures using phenix.refine (https://www.phenix-online.org/documentation/reference/refinement.html). This was done using phenix version 1.17.1-3660. We performed anisotropic refinement on all pairs where both PDBs had a resolution better than 1.5Å. All other refinement was run isotropically. Refinement used the following parameters:

- Refine strategy: individual sites + individual adp + occupancies
- Number of macro cycles: 8
- NQH flips: True
- Optimize xyz weight: True
- Optimize adp weight: True
- Hydrogen refine: Riding

We removed 102 structures because of incompatibility with our re-refinement pipeline due to breaks in the protein chain or ligand incompatibility. We removed 88 structures where the R-free increased by >2.5% compared to the value reported in the PDB header (**Supplementary Table 1; Supplementary Figure 2A**).

### Running qFit

qFit-3.0(Riley et al., 2021) (version 3.2.0) was run using a composite omit map and the re-refined structure on the default parameters(https://github.com/ExcitedStates/qfit-3.0/). We ran qFit on Amazon Web Services (AWS). We used an auto scaling cluster of images controlled by the scheduler via ParallelCluser. Please see the qFit github for a script that will install qFit on AWS’s default OS image, using conda to install its dependencies.

After qFit, we re-ran refinement as suggested by qFit-3.0. Briefly, this involves three rounds of refinement. The first refines coordinates only, the second goes through a cyclical round of refinement until the majority of low occupancy conformers are removed, and the third refinement polishes the structure, including hydrogens. The script used for post qFit refinement can be found here: https://github.com/ExcitedStates/qfit-3.0/blob/master/scripts/post/qfit_final_refine_xray.sh. We removed 100 structures because of incompatibility with refinement after qFit rebuilding.

### Quality Control

From our original dataset (n=1,205 pairs), we removed 28 apo structures that had a crystallographic additive or amino acid in the binding site that partially overlaid with the holo structure. We set a minimum ligand occupancy threshold of 0.15, which did not remove any pairs from our dataset. Chains were renamed according to the difference in distance between the two chains. We also re-numbered each chain based on the apo structure. We then superimposed the two structures using the pymol align function. We measured the alpha carbon root mean squared difference (RMSD) between the two structures as well as the difference in just binding site residues. Structures were removed if the mean RMSD of the entire structure was greater than 1Å or if the mean RMSD in the binding site residues were greater than 0.5Å. We removed two pairs based on these RMSD criteria.

We also assessed the difference in R-free values for each refinement step (before/after qFit). If the post refinement R-free value was 2.5% larger than the pre refinement R-value, then the structure was removed (n=85, 77 structures removed; **Supplementary Figure 2A**,**B**). Additionally, we compared the final R-free values between apo and holo pairs, removing pairs with R-free values with more than a 5% difference (n=16 pairs removed; **Supplementary Figure 2C**). We ran the clashscore function out of Molprobity(Williams et al., 2018) to identify severe clashes in our dataset. We removed any structures with a clashscore greater than 15, removing 52 structures. After filtering out pairs that failed our quality checks, our dataset contained 743 matched apo/holo pairs.

### Alternative Conformations

Side chains were considered alternative conformers if there was at least one atom that was modeled with an alternative conformer. Our re-refinement procedure changes the occupancy, coordinates, and B-factors of these conformations, but it does not add or delete conformations.

### B-factors

B-factors were assessed on a residue basis by averaging the B-factors of all heavy atoms for each residue. For residues with multiple conformations, we took the mean B-factor for all heavy atoms in each side chain, weighted by occupancy. For structures modeled anisotropically, we used the isotropic equivalent B-factor from phenix.

### Root Mean Squared Fluctuation (RMSF)

RMSF was chosen over root mean squared deviation as many alternative conformers were predicted to have the same occupancy, thus making it difficult to define which was the main conformer. RMSF was measured for each residue based on all side chain heavy atoms. RMSF finds the geometric center of each atom in all alternative conformers. It then takes the distance between the geometric mean of each conformer’s side chain heavy atoms and the overall geometric center. It then takes the squared mean of all of those distances, weighted by occupancy.

### Order Parameters

Order parameters were measured for each residue (except proline and glycine) by taking into account both the angle of alternative conformers (s2angle), by measuring the chi1 angle, and the B-factors of alpha or beta carbons along with an attached hydrogen(s2ortho)(Fenwick et al., 2014). To account for differences in B-factors as resolution changes, we investigated the correlation between order parameters in 32 apo lysozyme structures ranging in resolution from 1.1 to 2Å. We optimized the s2ortho parameter by looking for the normalization that would provide a slope closest to one and have the smallest root mean squared error. We normalized the s2ortho portion using the following equation:

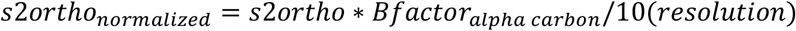

The final order parameter reported in the paper is:

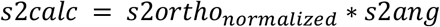

### Rotamer Analysis

Rotamers were determined using phenix.rotalyze(Williams et al., 2018) with manually relaxing the outlier criteria to 0.1%. Each alternative conformation has its own rotamer state. Rotamers were compared on a residue by residue basis between the holo and apo, taking into account each rotamer state for each alternative conformation. Residues were classified as “no change” if rotamers matched across the apo and holo residue, “distinct” if the matched residue shared no rotamer assignments. Residues were classified as “remodeled-holo loss” if distinct, additional rotameric conformations were populated in the apo residue only, and “remodeled - holo gain” if distinct, additional rotameric conformations were populated in the holo residue only.

### Solvent Exposed Surface Area

We calculated the relative accessible surface area (RASA) using Define Secondary Structure of Proteins (DSSP)(Kabsch and Sander, 1983) with the Tien et al (Tien et al., 2013) definition of Max accessible surface area (MaxASA). Residues with a RASA of ≥20% were considered solvent exposed.

### Ligand Analysis

We obtained the ligand properties using RDkit (version 2021.03.2) by importing SDF files of each ligand in our dataset. To account from the multiple hypothesis testing, we applied a Bonferroni correction, with an alpha of 0.05, as we were testing 10 hypotheses leaving us with a corrected significance value of 0.005.

Occupancy of the ligands were taken directly from the PDB file and correspond with the ligand occupancy from the deposited structure. Ligand B-factors were normalized using the mean alpha carbon B-factor of all residues in the structure.

If there were multiple ligands of interest in a structure, we looked at the properties of the ligand and surrounding protein residues in chain A or in the lowest alphabetical chain.

### Protein Type Analysis

Protein names and enzyme names were extracted from Uniprot(UniProt Consortium, 2015). Names and properties were connected using PDB IDs.

### Statistics

Paired Wilcoxen test was used for all matched properties (comparing holo v. apo matched residues or structures). Individual Mann-Whitney U test was used for all non-match properties, including ligand properties. Two-sided t-test was used to compare the significance of the slopes.

### Code

Code can be found in the following repositories:

- Dataset selection: https://github.com/stephaniewankowicz/PDB_selection_pipeline
- Refinement/qFit pipeline: https://github.com/stephaniewankowicz/refinement_qFit
- Analysis/Figures: https://github.com/fraser-lab/Apo_Holo_Analysis
- qFit: https://github.com/ExcitedStates/qfit-3.0.

## Supplementary Tables

**Supplementary Table 1 (Crystallographic Additives)**

**Supplementary Table 2 (Holo/Apo Pairs)**

**Supplementary Table 3 (Apo/Apo Pairs)**

## Acknowledgements

We thank James Holton, Tony Capra, and Dan Herschlag for helpful comments and suggestions. JSF is supported by NIH GM123159 and GM124149, and a Sanghvi-Agarwal Innovation Award. SAW is supported by NSF GRFP 2034836.

## Competing interests

SHPdO and HvdB are employees of Atomwise Inc. JSF has equity, has received consulting fees, and has sponsored research agreements with Relay Therapeutics.

## REFERENCES

van den Bedem, H., and Fraser, J.S. (2015). Integrative, dynamic structural biology at atomic resolution--it’s about time. Nat. Methods 12, 307–318.

van den Bedem, H., Dhanik, A., Latombe, J.C., and Deacon, A.M. (2009). Modeling discrete heterogeneity in X-ray diffraction data by fitting multi-conformers. Acta Crystallogr. D Biol. Crystallogr. 65, 1107–1117.

Berman, H. (2002). Protein Data Bank Project at Rutgers University.

Berman, H.M., Westbrook, J., Feng, Z., Gilliland, G., Bhat, T.N., Weissig, H., Shindyalov, I.N., and Bourne, P.E. (2000). The Protein Data Bank. Nucleic Acids Res. 28, 235–242.

Bhat, T.N. (1989). Correlation between occupancy and temperature factors of solvent molecules in crystal structures of proteins. Acta Crystallogr. A 45 (Pt 1), 145–146.

Boehr, D.D., Nussinov, R., and Wright, P.E. (2009). The role of dynamic conformational ensembles in biomolecular recognition. Nat. Chem. Biol. 5, 789–796.

Breiten, B., Lockett, M.R., Sherman, W., Fujita, S., Al-Sayah, M., Lange, H., Bowers, C.M., Heroux, A., Krilov, G., and Whitesides, G.M. (2013). Water networks contribute to enthalpy/entropy compensation in protein-ligand binding. J. Am. Chem. Soc. 135, 15579–15584.

Bromberg, S., and Dill, K.A. (1994). Side-chain entropy and packing in proteins. Protein Sci. 3, 997–1009.

Burnley, B.T., Afonine, P.V., Adams, P.D., and Gros, P. (2012). Modelling dynamics in protein crystal structures by ensemble refinement. Elife 1, e00311.

Caldararu, O., Ekberg, V., Logan, D.T., Oksanen, E., and Ryde, U. (2021). Exploring ligand dynamics in protein crystal structures with ensemble refinement. Acta Crystallogr D Struct Biol 77, 1099–1115.

Caro, J.A., Harpole, K.W., Kasinath, V., Lim, J., Granja, J., Valentine, K.G., Sharp, K.A., and Wand, A.J. (2017). Entropy in molecular recognition by proteins. Proc. Natl. Acad. Sci. U. S. A. 114, 6563–6568.

Caro, J.A., Valentine, K.G., and Joshua Wand, A. (2021). Structural origins of protein conformational entropy.

Carugo, O. (1999). Correlation between occupancy and B factor of water molecules in protein crystal structures. Protein Eng. 12, 1021–1024.

Cheng, Y., Grigorieff, N., Penczek, P.A., and Walz, T. (2015). A primer to single-particle cryo-electron microscopy. Cell 161, 438–449.

Doig, A.J., and Sternberg, M.J. (1995). Side-chain conformational entropy in protein folding. Protein Sci. 4, 2247–2251.

Eshun-Wilson, L., Zhang, R., Portran, D., Nachury, M.V., Toso, D.B., Löhr, T., Vendruscolo, M., Bonomi, M., Fraser, J.S., and Nogales, E. (2019). Effects of α-tubulin acetylation on microtubule structure and stability. Proc. Natl. Acad. Sci. U. S. A. 116, 10366–10371.

Fenwick, R.B., van den Bedem, H., Fraser, J.S., and Wright, P.E. (2014). Integrated description of protein dynamics from room-temperature X-ray crystallography and NMR. Proc. Natl. Acad. Sci. U. S. A. 111, E445–E454.

Fleck, M., Polyansky, A. A., & Zagrovic, B. (2018) Self-Consistent Framework Connecting Experimental Proxies of Protein Dynamics with Conformational Entropy. Journal of chemical therory and computation, 14(7), 3796–3810.

Fraser, J.S., van den Bedem, H., Samelson, A.J., Lang, P.T., Holton, J.M., Echols, N., and Alber, T. (2011). Accessing protein conformational ensembles using room-temperature X-ray crystallography. Proc. Natl. Acad. Sci. U. S. A. 108, 16247–16252.

Frederick, K.K., Marlow, M.S., Valentine, K.G., and Wand, A.J. (2007). Conformational entropy in molecular recognition by proteins. Nature 448, 325–329.

Gagné, S. M., Tsuda, S., Spyracopoulos, L., Kay, L. E., & Sykes, B. D. (1998). Backbone and methyl dynamics of the regulatory domain of troponin C: anisotropic rotational diffusion and contribution of conformational entropy to calcium affinity. Journal of molecular biology, 278(3), 667–686.

Gaudreault, F., Chartier, M., and Najmanovich, R. (2012). Side-chain rotamer changes upon ligand binding: common, crucial, correlate with entropy and rearrange hydrogen bonding. Bioinformatics 28, i423–i430.

Gohlke, H., Kuhn, L.A., and Case, D.A. (2004). Change in protein flexibility upon complex formation: analysis of Ras-Raf using molecular dynamics and a molecular framework approach. Proteins 56, 322–337.

Gutteridge, A., and Thornton, J. (2005). Conformational changes observed in enzyme crystal structures upon substrate binding. J. Mol. Biol. 346, 21–28.

Hoffmann, F., Mulder, F.A., & Schäfer, L.V. (2021). How much entropy is contained in NMR relaxation parameters?

Jain, A.N., Cleves, A.E., Brueckner, A.C., Lesburg, C.A., Deng, Q., Sherer, E.C., and Reibarkh, M.Y. (2020). XGen: Real-Space Fitting of Complex Ligand Conformational Ensembles to X-ray Electron Density Maps. J. Med. Chem. 63, 10509–10528.

Jhaveri, K., Ha, B.R., Yap, T.A., Hamilton, E., Rugo, H.S., Goldman, J.W., Dann, S., Liu, F., Wong, G.Y., Krupka, H., et al. (2021). The evolution of cyclin dependent kinase inhibitors in the treatment of cancer. Expert Rev. Anticancer Ther.

Kabsch, W., and Sander, C. (1983). Dictionary of protein secondary structure: pattern recognition of hydrogen-bonded and geometrical features. Biopolymers 22, 2577–2637.

Keedy, D.A., Fraser, J.S., and van den Bedem, H. (2015). Exposing Hidden Alternative Backbone Conformations in X-ray Crystallography Using qFit. PLoS Comput. Biol. 11, e1004507.

Kim, J., Ahuja, L.G., Chao, F.-A., Xia, Y., McClendon, C.L., Kornev, A.P., Taylor, S.S., and Veglia, G. (2017). A dynamic hydrophobic core orchestrates allostery in protein kinases. Sci Adv 3, e1600663.

Kuriyan, J., Petsko, G.A., Levy, R.M., and Karplus, M. (1986). Effect of anisotropy and anharmonicity on protein crystallographic refinement. An evaluation by molecular dynamics. J. Mol. Biol. 190, 227–254.

Kuzmanic, A., Pannu, N.S., and Zagrovic, B. (2014). X-ray refinement significantly underestimates the level of microscopic heterogeneity in biomolecular crystals. Nat. Commun. 5, 3220.

Lang, P.T., Ng, H.-L., Fraser, J.S., Corn, J.E., Echols, N., Sales, M., Holton, J.M., and Alber, T. (2010). Automated electron-density sampling reveals widespread conformational polymorphism in proteins. Protein Sci. 19, 1420–1431.

Liebschner, D., Afonine, P.V., Baker, M.L., Bunkóczi, G., Chen, V.B., Croll, T.I., Hintze, B., Hung, L.W., Jain, S., McCoy, A.J., et al. (2019). Macromolecular structure determination using X-rays, neutrons and electrons: recent developments in Phenix. Acta Crystallogr D Struct Biol 75, 861–877.

Mobley, D.L., and Dill, K.A. (2009). Binding of small-molecule ligands to proteins: “what you see” is not always “what you get.” Structure 17, 489–498.

Moorman, V.R., Valentine, K.G., and Wand, A.J. (2012). The dynamical response of hen egg white lysozyme to the binding of a carbohydrate ligand. Protein Sci. 21, 1066–1073.

Olsson, T.S.G., Williams, M.A., Pitt, W.R., and Ladbury, J.E. (2008). The thermodynamics of protein-ligand interaction and solvation: insights for ligand design. J. Mol. Biol. 384, 1002–1017.

Riley, B.T., Wankowicz, S.A., de Oliveira, S.H.P., van Zundert, G.C.P., Hogan, D.W., Fraser, J.S., Keedy, D.A., and van den Bedem, H. (2021). qFit 3: Protein and ligand multiconformer modeling for X-ray crystallographic and single-particle cryo-EM density maps. Protein Sci. 30, 270–285.

Rosenbaum, D., Garnelo, M., Zielinski, M., Beattie, C., Clancy, E., Huber, A., Kohli, P., Senior, A.W., Jumper, J., Doersch, C., et al. (2021). Inferring a Continuous Distribution of Atom Coordinates from Cryo-EM Images using VAEs.

Steuber, H., Zentgraf, M., Gerlach, C., Sotriffer, C.A., Heine, A., and Klebe, G. (2006). Expect the Unexpected or Caveat for Drug Designers: Multiple Structure Determinations Using Aldose Reductase Crystals Treated under Varying Soaking and Co-crystallisation Conditions. Journal of Molecular Biology 363, 174–187.

Tien, M.Z., Meyer, A.G., Sydykova, D.K., Spielman, S.J., and Wilke, C.O. (2013). Maximum allowed solvent accessibilites of residues in proteins. PLoS One 8, e80635.

Tzeng, S.-R., and Kalodimos, C.G. (2012). Protein activity regulation by conformational entropy. Nature488, 236–240.

UniProt Consortium (2015). UniProt: a hub for protein information. Nucleic Acids Res. 43, D204–D212.

Verteramo, M.L., Stenström, O., Ignjatović, M.M., Caldararu, O., Olsson, M.A., Manzoni, F., Leffler, H., Oksanen, E., Logan, D.T., Nilsson, U.J., et al. (2019). Interplay between Conformational Entropy and Solvation Entropy in Protein–Ligand Binding. Journal of the American Chemical Society 141, 2012–2026.

Wand, A.J., and Sharp, K.A. (2018). Measuring Entropy in Molecular Recognition by Proteins. Annu. Rev. Biophys. 47, 41–61.

Wang, Y., Manu, V.S., Kim, J., Li, G., Ahuja, L.G., Aoto, P., Taylor, S.S., and Veglia, G. (2019). Globally correlated conformational entropy underlies positive and negative cooperativity in a kinase’s enzymatic cycle. Nature Communications 10.

Wicker, J.G.P., and Cooper, R.I. (2015). Will it crystallise? Predicting crystallinity of molecular materials. CrystEngComm 17, 1927–1934.

Williams, C.J., Headd, J.J., Moriarty, N.W., Prisant, M.G., Videau, L.L., Deis, L.N., Verma, V., Keedy, D.A., Hintze, B.J., Chen, V.B., et al. (2018). MolProbity: More and better reference data for improved all-atom structure validation. Protein Sci. 27, 293–315.

Woldeyes, R.A., Sivak, D.A., and Fraser, J.S. (2014). E pluribus unum, no more: from one crystal, many conformations. Curr. Opin. Struct. Biol. 28, 56–62.

Wong, K.-B., and Daggett, V. (1998). Barstar Has a Highly Dynamic Hydrophobic Core: Evidence from Molecular Dynamics Simulations and Nuclear Magnetic Resonance Relaxation Data†. Biochemistry 37, 11182–11192.

Wood, D.J., and Endicott, J.A. (2018). Structural insights into the functional diversity of the CDK-cyclin family. Open Biol. 8.

Zhou, H.-X., and Gilson, M.K. (2009). Theory of free energy and entropy in noncovalent binding. Chem. Rev. 109, 4092–4107.

van Zundert, G.C.P., Hudson, B.M., de Oliveira, S.H.P., Keedy, D.A., Fonseca, R., Heliou, A., Suresh, P., Borrelli, K., Day, T., Fraser, J.S., et al. (2018). qFit-ligand Reveals Widespread Conformational Heterogeneity of Drug-Like Molecules in X-Ray Electron Density Maps. J. Med. Chem. 61, 11183–11198.

