## Supplementary Figures for "Ligand binding remodels protein side chain conformational heterogeneity"

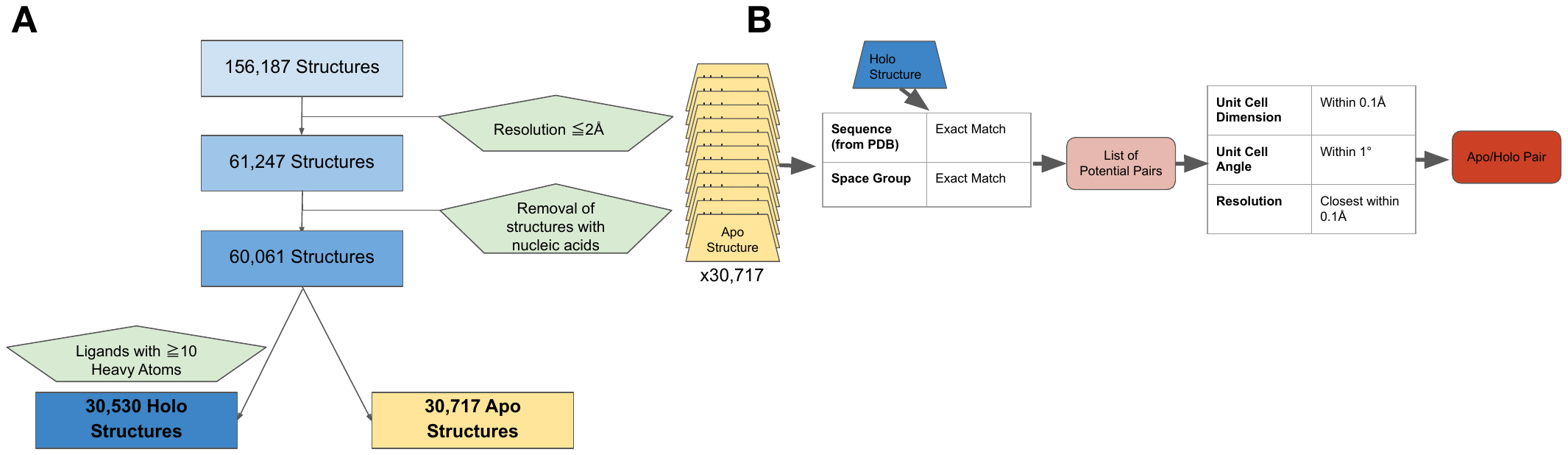

**Supplementary Figure 1***. (A) To select holo/apo matched pairs, we first categorized the PDB structures into holo or apo structures, removing structures with a resolution worse than 2Å, not resolved using X-ray crystallography, and those that include nucleic acids. Holo structures (n=30,530) were required to have a ligand, not including common crystallographic additives, with 10 or more heavy atoms. All others were classified as apo (n=30,171). (B) For every holo structure, we compared it to the 30,717 apo structures first matching for exact sequence and space group and controlling for similar unit cell dimensions (within 0.1 Å) and angles (within 1 degree). Finally, we selected the structures paired for resolution within 0.1 Å.*

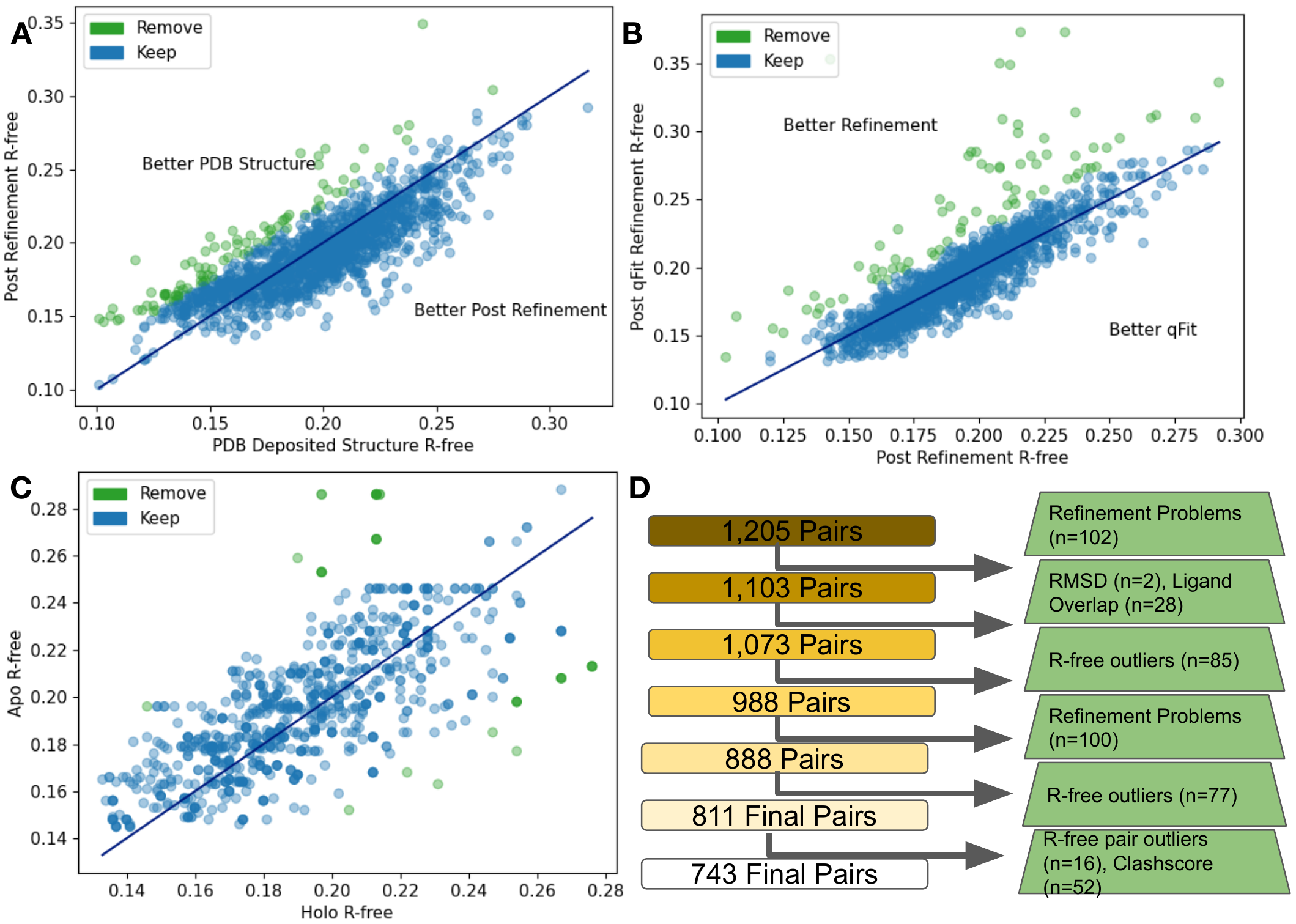

**Supplementary Figure 2***. (A) The differences in R-free values between the PDB deposited structures and after re-refinement. 85 structures were removed (green) as their R-free increased by more than 2.5%. (B) The difference in R-free statistics between the re-refined structures and the qFit structures. 77 structures were removed (green) as their R-free increased by more than 2.5%. (C) The difference in R-free statistics in qFit structures between the holo and apo structure. 16 pairs were removed (green) as ther R-free statistics differed by 5% or more between the pairs. (D) Flowchart representing our quality control process, with removed structures in green boxes.*

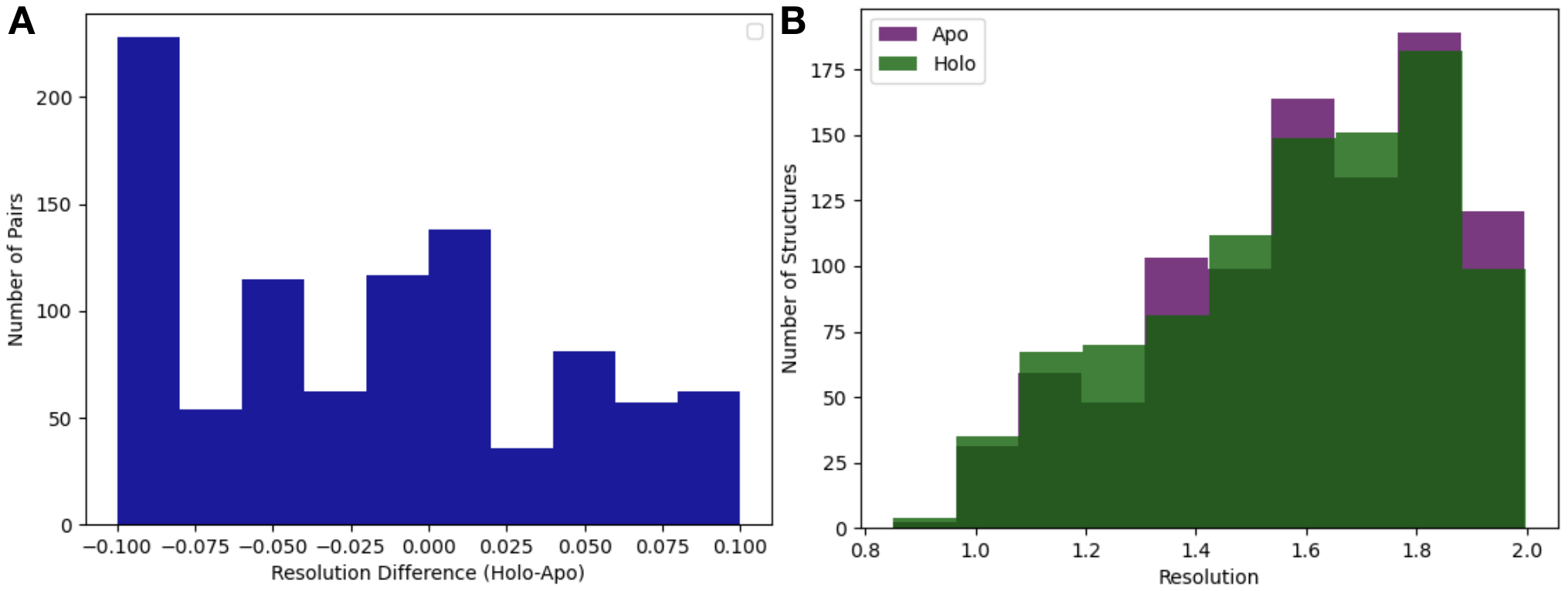

**Supplementary Figure 3***. (A) Resolution difference between pairs (holo-apo). The median pairwise difference was 0.01*Å, with slightly better resolution in the apo structures, *and the standard deviation was 0.06*Å*. (B) The distribution of resolution (median=1.6*Å*) of the apo (n=432) and holo (n=743) dataset. The median apo resolution was 1.58*Å *and the median holo resolution was 1.58*Å*.*

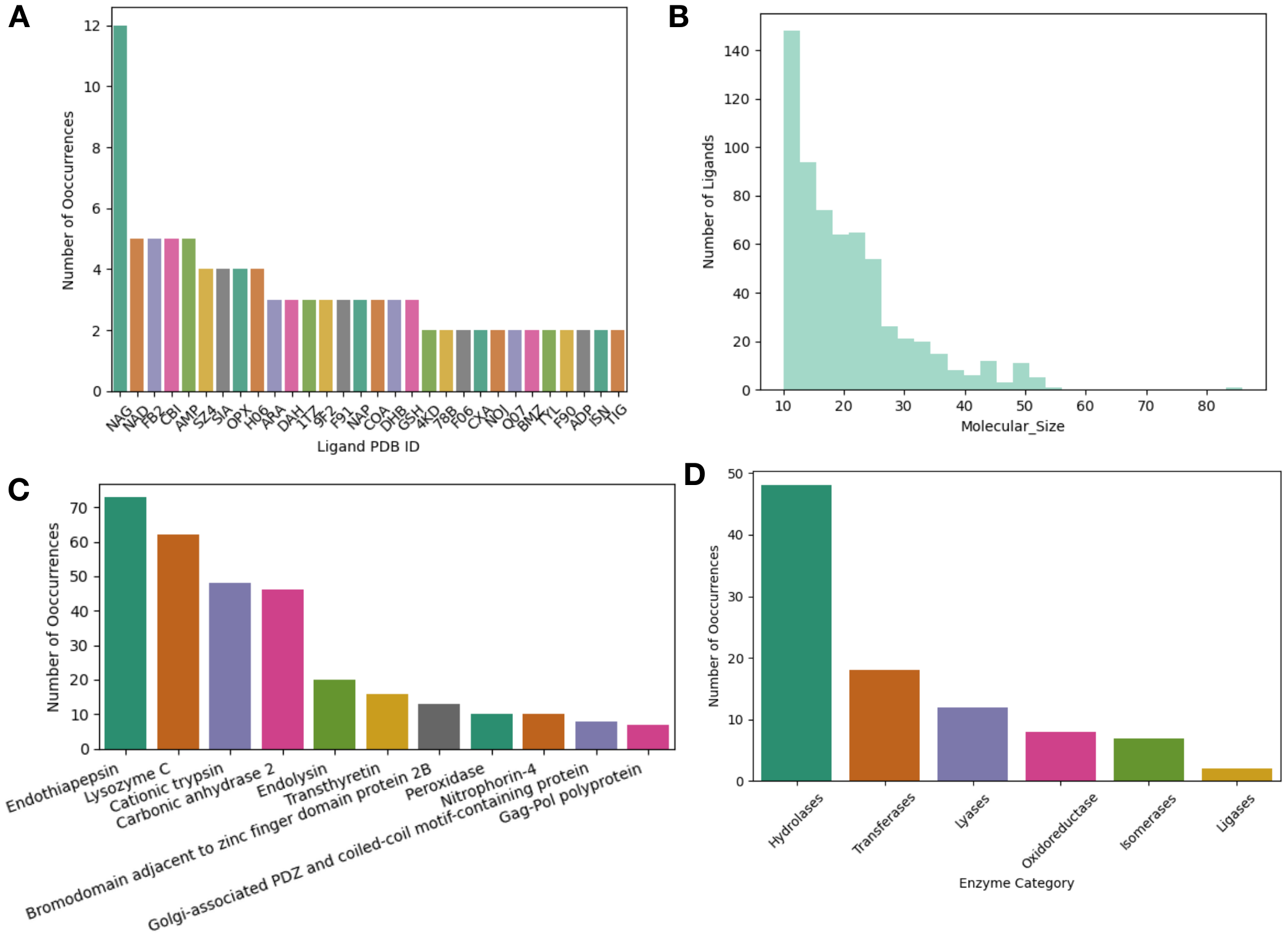

**Supplementary Figure 4***. (A)The top 30 ligands in our dataset by PDB chemical ID. NAG (2-acetamido-2-deoxy-beta-D-glucopyranose) and H06 ((E)-4-((2-nicotinoylhydrazono)methyl) benzimidamide) where the most frequent ligands in our dataset. (B) The distribution of the number of heavy atoms of a ligand of interest. The median number of heavy atoms was 19. There were only 10 very large ligands (>50 heavy atoms, e.g. Atazanavir). (C) The most common proteins in our dataset. Eleven proteins in our dataset were included in 6 or more pairs. This included our most common proteins including: Endothiopepsin (n=73 pairs), Lysozyme (n=62 pairs), Trypsin (n=48 pairs), and Carbonic Anhydrase 2 (n=46 pairs). (D) The distribution of enzymes (n=95) based on their Enzyme Commission Number.*

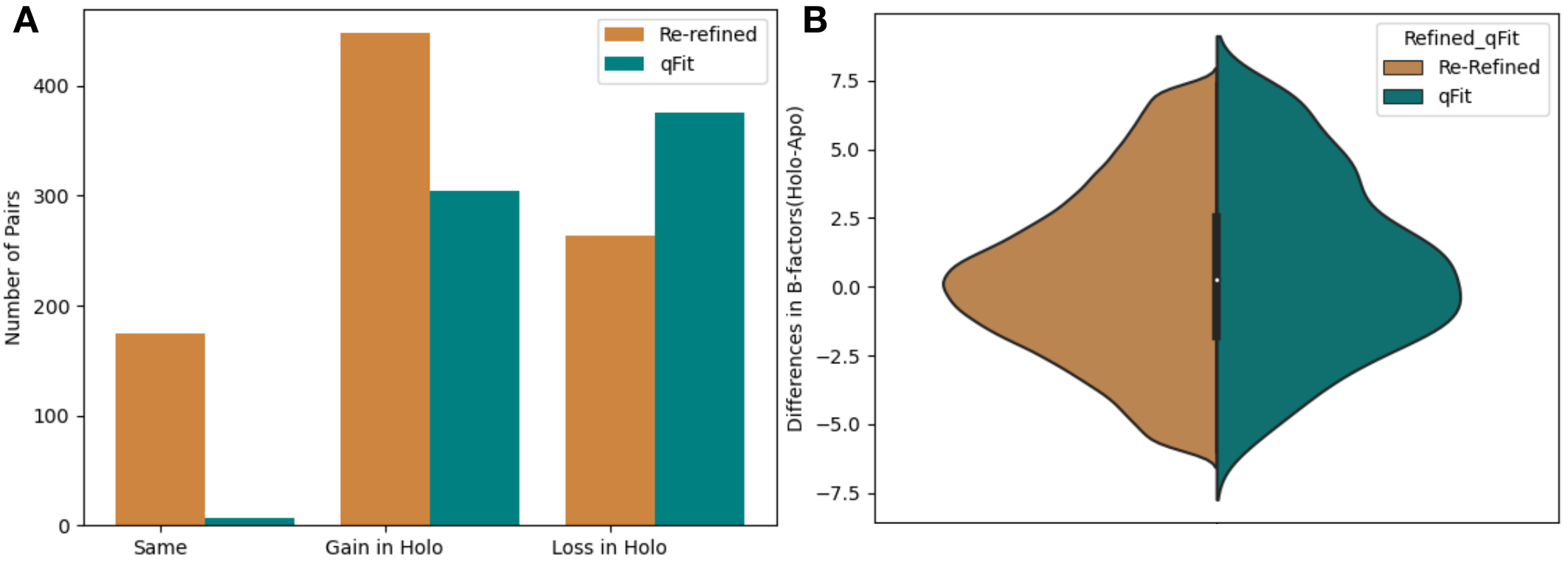

**Supplementary Figure 5****.** (A) *The change in the number of alternative conformers (holo-apo) across all residues. In the re-refined dataset (orange), the majority models have a gain of the number of alternative conformers in the holo, with the second most common category being a loss of alternative conformers. In the qfit dataset (teal), the majority of structures lose an alternative conformer in the holo model, with the second most common category being gaining an alternative conformer. (B) The difference in B-factors across all residues. There was a slight increase in B-factors in holo modelss in both the re-refined and the qFit datasets.*

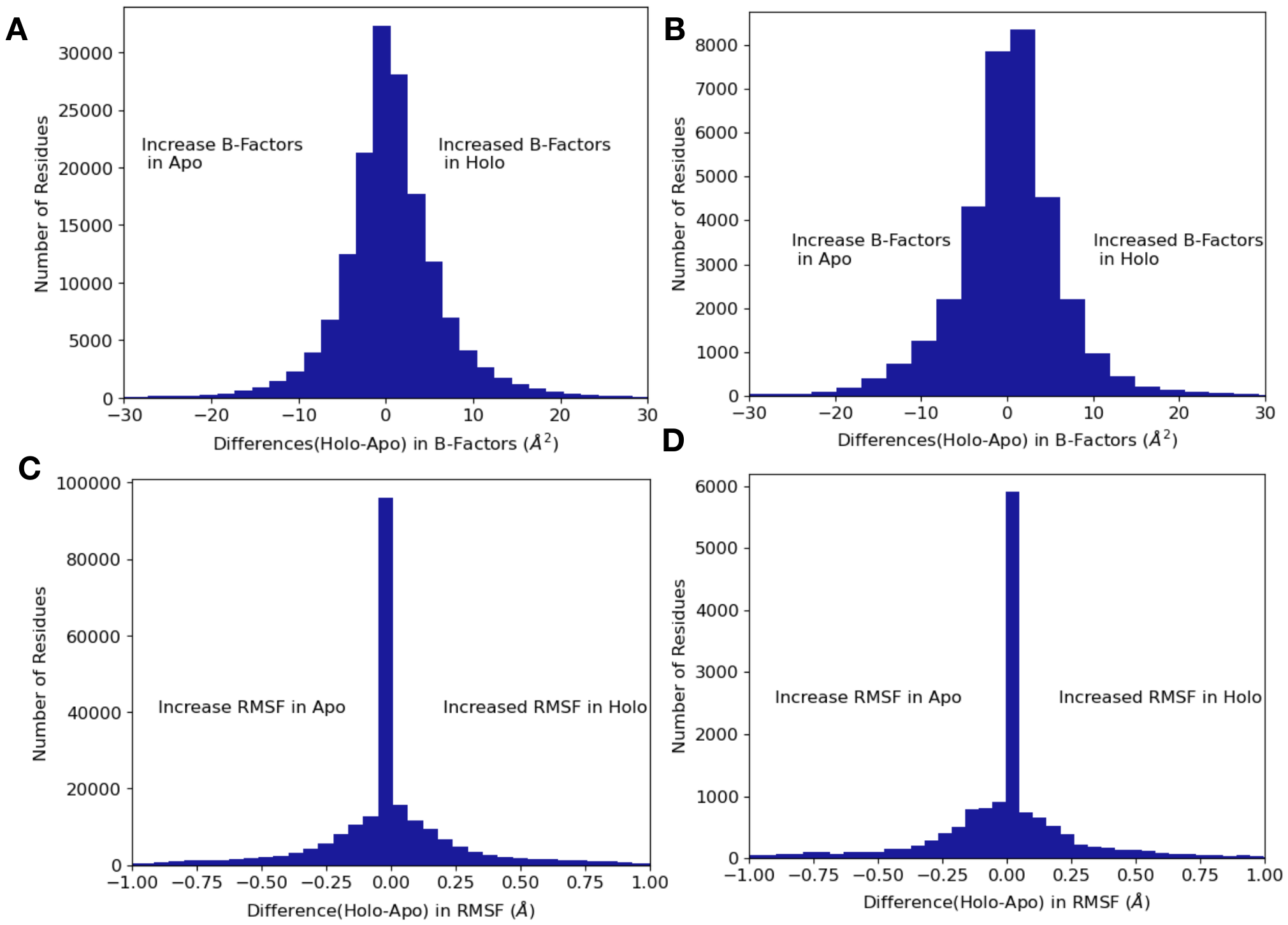

**Supplementary Figure 6***. (A) The difference in B-factors between holo and apo pairs. The range of the difference in B-factors was -199.8*Å^2^ *to 197.0*Å^2^*, here we remove the most 10% extreme values, which are due to poor density in loop regions leading to high B-factors for those individual residues. Across all residues, on average B-factors were higher in holo structures compared to apo (*0.34Å^2^, median difference (holo-apo); p=4.4x10^-208^, Wilcoxon signed rank test*). (B) In binding site residues, B-factors were on average the same between holo and apo residues (*0.06Å^2^, median difference in B-factors; p=0.7, Wilcoxon signed rank test). *(C) Across all residues, apo residues had a higher RMSF compared to holo residues (0.17*Å vs. *0.16*Å, mean RMSF; *-0.006, mean difference: p=4.5x10^-29^ ,Wilcoxon signed rank test). (D) Within binding site residues, apo residues also had a higher RMSF, compared to holo residues (0.17*Å vs. *0.15*Å, mean RMSF*; -0.02, mean difference; p=3.7x10^-8^, Wilcoxon signed rank test).*

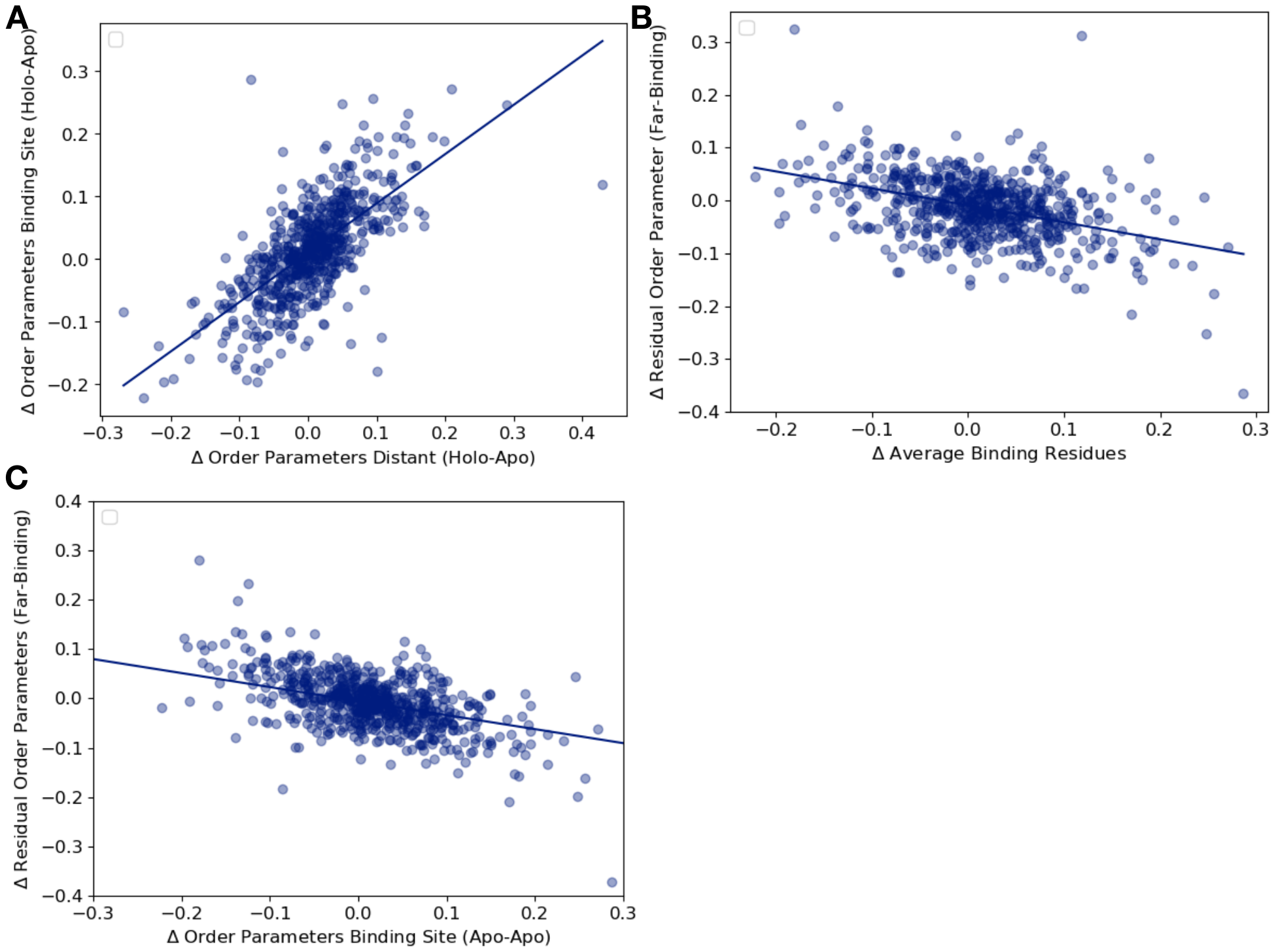

**Supplementary Figure 7.** *(A) The relationship between the average order parameter in distant, non-solvent exposed residues versus the average order parameters in binding site residues (n=743, slope=0.79, r^2^=0.65; p=6.5x10^-89^, two-sided t-test). (B) The relationship between the residual order parameters in all distant residues versus binding site residue order parameters (n=743, slope=-0.32, r^2^=0.17; p=4.6x10^-28^, two-sided t-test). (C) The relationship between the residual order parameters in distant, non-solvent exposed residues versus binding site residues in the apo and apo control dataset residues (n=283, slope=-0.28, r^2^=0.20; p=1.8x10^-34^, two-sided t-test).*

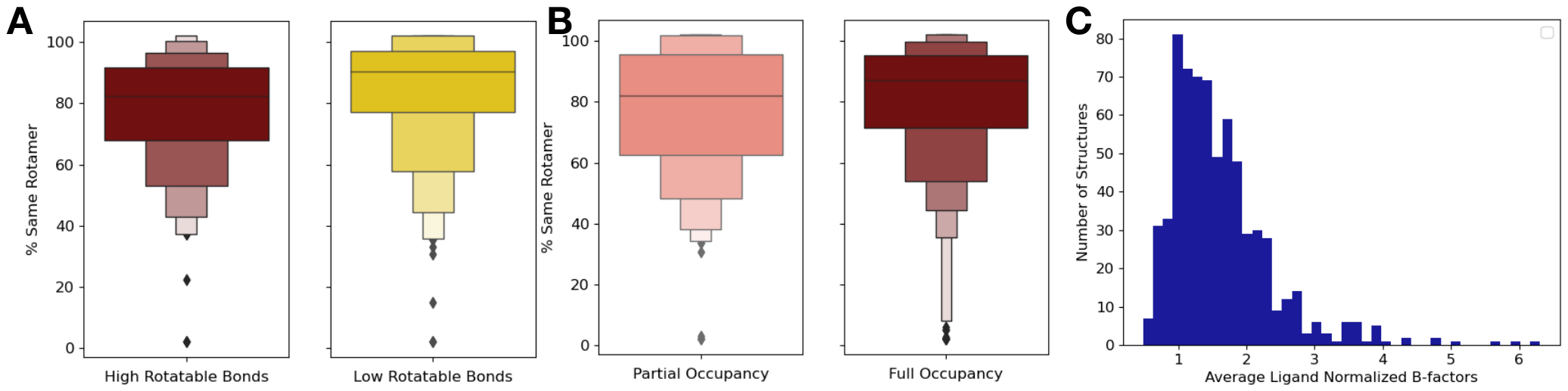
***Supplementary Figure 8.*** *(A) We explored if the top and bottom quartiles of rotatable bond ligands were associated with an increase or decrease of rotamer changes, as defined as the percentage of close residues with the same rotamer in the holo and apo structure. The ligands in the top quartile of rotatable bonds had less rotamers that were the same between holo and apo structures versus ligands in the bottom quartile of rotatable bonds (80% vs. 88%, median same percentage of rotamers, p=0.001, independent Mann Whitney U test). (B) There was no significant difference in the percentage of the same rotamers between partially occupied and fully occupied ligands (80% vs. 85%, median percentage of the same rotamer; p=0.11, independent Mann Whitney U test). (C) In fully occupied ligands, the median B-factor was 24.8, with a range of 5.5 to 99.3.*

**Supplementary Figure 9****.**  *We looked at the difference in order parameters (holo-apo) and the supporting density for specific residues between the apo (PDB: 1PW2, purple), type II (PDB: 1PXI, 3QQL, teal), and type I (PDB: 2A0C, 3QTW, 3R1Q, salmon) inhibitors. All density is shown at 1 sigma. (A) Valine 18, one of the ligand contacts for both the type I and type II inhibitors. Across all holo structures this residue becomes more rigid, including losing an alternative conformer and changing rotamers in the holo structure. This residue is also a part of the blue cluster in the heatmap. (B) Glutamine 127, one of the ligand contacts for both type I and type II inhibitors. This residue has two very different alternative conformers in the apo structure. In the type II inhibitor structure, there are again two very different alternative conformers whereas in the type I inhibitor structure, there are three very similar alternative conformers. This residue is also a part of the blue cluster in the heatmap. (C) Tyrosine 15 in the P-loop has varying differences in order parameters. In the type II inhibitor, this tyrosine gets more rigid, along with the rest of the p-loop, however in the type I inhibitor structures, this tyrosine along with the rest of the p-loop becomes more dynamic.*
